# Frontal Cortex Gates Distractor Stimulus Encoding in Sensory Cortex

**DOI:** 10.1101/2022.03.31.486430

**Authors:** Zhaoran Zhang, Edward Zagha

## Abstract

Frontal cortex suppresses behavioral responses to distractor stimuli. One possible mechanism by which this occurs is by modulating sensory responses in sensory cortex. However, it is currently unknown how frontal cortex modulations of sensory cortex contribute to distractor response suppression. We trained mice to respond to target stimuli in one whisker field and ignore distractor stimuli in the opposite whisker field. During expert task performance, optogenetic inhibition of frontal cortex increased behavioral responses to distractor stimuli. During expert task performance, within sensory cortex we observed expanded propagation of target stimulus responses and contracted propagation of distractor stimulus responses. In contrast to current models of frontal cortex function, frontal cortex did not substantially modulate the response amplitude of preferred stimuli. Rather, frontal cortex specifically suppressed the propagation of distractor stimulus responses, thereby preventing target-preferring neurons from being activated by distractor stimuli. Single unit analyses revealed that wMC decorrelates target and distractor stimulus encoding in target-preferring S1 neurons, which likely improves selective target stimulus detection by downstream readers. Moreover, we observed proactive top-down modulation from frontal to sensory cortex, through the preferential activation of GABAergic neurons. Overall, our study provides important mechanistic details about how frontal cortex gates sensory propagation in sensory cortex to prevent behavioral responses to distractor stimuli.

**Highlights:**

- Pairing of frontal cortex optogenetic inhibition with sensory cortex recordings during a target-distractor Go/NoGo task.
- During expert task performance, we observed target stimulus response expansion and distractor stimulus response contraction.
- Optogenetic inhibition of frontal cortex increased false alarm rates and selectively increased the propagation of distractor evoked responses into target-aligned sensory cortex.
- Even before stimulus onset, frontal cortex preferentially drives GABAergic neurons in distractor-aligned sensory cortex.

## Introduction

To achieve our goals, we must respond to task-relevant target stimuli and not respond to task-irrelevant distractors. The ability to suppress behavioral responses to distractor stimuli is a component of impulse control, which is a core cognitive process. Impairments in distractor response suppression underlie some of the behavioral and cognitive dysfunctions in neuropsychiatric disorders such as attention-deficit/hyperactivity disorder (ADHD), obsessive– compulsive disorder, and substance abuse disorder (Chamberlain & Sahakian, 2007; Hester & Garavan, 2004).

Frontal cortex is an essential mediator of distractor response suppression. Wide-ranging studies across multiple species and sensory modalities have shown that disruptions of frontal cortex impair the ability to suppress behavioral responses to distractor stimuli (Buia & Tiesinga, 2008; Bussey et al., 1997; Eichenbaum et al., 1974; Eichenbaum et al., 1983; Hvoslef-Eide et al., 2018; Iversen & Mishkin, 1970; Kolb et al., 1974; Konorski, 1972; Leimkuhler & Mesulam, 1985; Li et al., 2020; Markowitsch & Pritzel, 1977; Picton et al., 2007; Swick et al., 2008; Salmaso & Denes, 1982; Zagha et al., 2015). In healthy human subjects, enhanced activation of the frontal cortex correlates with enhanced distractor response suppression (Carter et al., 1998; Li et al., 2006; Rubia et al., 2001). However, the mechanisms by which frontal cortex mediates distractor response suppression are poorly understood, due to the lack of studies combining causal manipulations of frontal cortex with sensory response recordings during target-distractor discrimination tasks.

Sensory responses are robustly modulated during performance in target-distractor discrimination tasks. The dominant findings are enhancements of target stimulus responses and suppressions of distractor stimulus responses (Anton-Erxleben et al., 2009; Atiani et al., 2009; Fritz et al., 2003; Fritz et al., 2005; Fritz et al., 2010; Liu et al., 2020; Moran & Desimone, 1985; Polley et al., 2006; David et al., 2012; Winkowski et al., 2018). However, these prior studies did not include causal manipulations of frontal cortex, and therefore it is unknown which of these modulations are driven by frontal cortex. Stimulation or suppression of frontal cortex does modulate sensory responses within sensory cortex (Angelucci et al., 2002; Hupé et al., 1998; Liu et al., 2020; Liu et al., 2021; Manita et al., 2015; Moore & Armstrong, 2003; Nassi et al., 2013; Noudoost & Moore, 2011; Winkowski et al., 2018; Zagha et al., 2013), which can impact behavior (Liu et al., 2021; Noudoost & Moore, 2011). Most studies of frontal cortex modulations of sensory processing emphasize enhancement or suppression of sensory cortex neurons to their preferred (optimal) stimuli, as opposed to the propagation of these signals within sensory cortex. And yet, these causal studies were not performed during target-distractor discrimination tasks, and therefore it is unknown which modulations contribute to distractor response suppression. Furthermore, it is generally assumed that frontal cortex modulates sensory responses through functional feedback/reactive signals (Angelucci et al., 2002; Hupé et al., 1998; Manita et al., 2015; Moore & Armstrong, 2003; Nassi et al., 2013). An alternative framework of proactive modulation (Zagha 2020) has yet to be causally tested.

In this study we combined frontal cortex suppression with sensory cortex single unit recordings in mice performing a target-distractor discrimination task. The task is a Go/No-Go operant task, in which mice learn to respond to rapid stimuli in one whisker field (target) and ignore identical stimuli in the opposite whisker field (distractor). Primary somatosensory cortex (S1) receives bottom-up inputs from contralateral whiskers. Additionally, S1 receives top-down inputs from ipsilateral frontal cortex, most robustly from the whisker region of motor cortex (wMC). Stimulation and suppression studies have demonstrated robust yet varied impacts of wMC activity on S1, including through excitation (Jung et al., 2022; Lee et al., 2008; Manita et al., 2015; Petreanu et al., 2012; Rocco & Brumberg, 2007; Xu et al., 2012; Zagha et al., 2013), inhibition (Kinnischtzke et al., 2014; Zagha et al., 2016), and dis-inhibition (Lee et al., 2013). The impacts of wMC on S1 in the context of distractor suppression is unknown.

## Results

### Performance in a selective detection task

Mice were trained to respond to target stimuli in one whisker field and ignore distractor stimuli in the opposite whisker field (Fig. 1 A,B,C). Given the lateralization of the mouse whisker representation, this task establishes target-aligned and distractor-aligned cortices that are symmetric across hemispheres and contralateral to the site of stimulus delivery (Fig. 1A). Performance in this task was quantified based on behavioral responses on target trials (hit rate), distractor trials (false alarm rate), and catch trials (spontaneous response rate). Mice were considered expert in this task once they achieved a discrimination d prime (d’, separation of hit rate and false alarm rate) greater than 1 for three consecutive days. Optogenetic perturbations and electrophysiological recordings were performed in expert mice while performing this selective detection task. Two stimulus amplitudes were applied in each session (equal for target and distractor trials): ‘large’ amplitude stimuli near the saturation of the hit rate psychometric curve and ‘small’ amplitude stimuli within the dynamic range. Unless otherwise indicated, our data analyses reference the large amplitude stimuli.

**Fig. 1:**
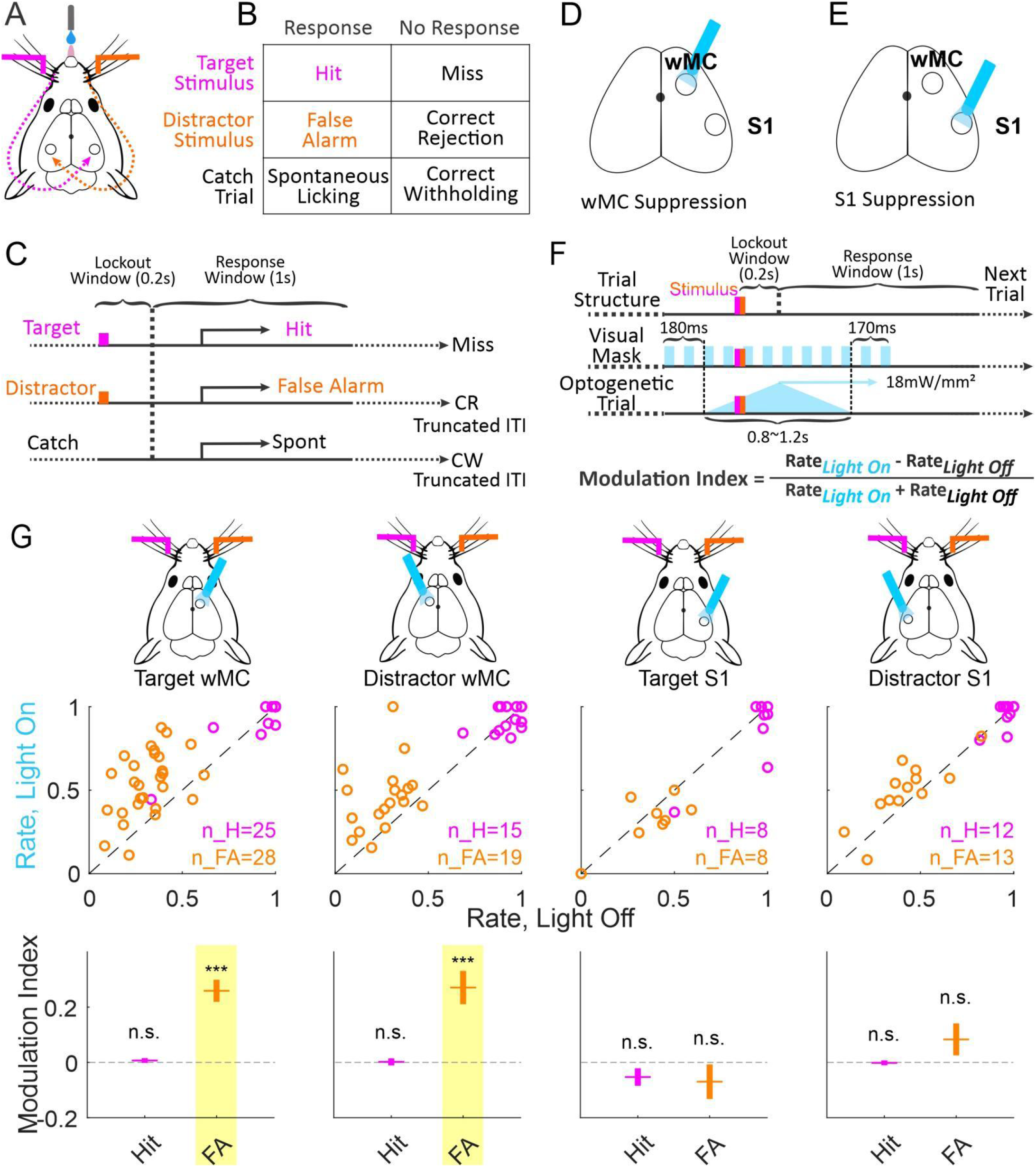
Suppressing either target-aligned or distractor-aligned wMC increases false alarm (FA) rates. (A) Illustration of the behavioral task setup. Mice are head-fixed in the behavioral rig with piezo-controlled paddles within their whisker fields bilaterally. Stimulus-triggered neural responses propagate to S1 in the contralateral hemisphere. (B) Task trial types and outcomes: for each trial there can be a target stimulus (hit or miss), a distractor stimulus (false alarm or correct rejection), or no stimulus catch trial (spontaneous licking or correct withholding). (C) Task structure: after whisker stimulus onset there is a 0.2s lockout window, followed by a 1s response window. Responding outside of the response window is punished by a time-out (resetting the inter-trial interval, ITI). Correct rejection and correct withhold are rewarded by a shortened ITI and subsequent target trial. Hits are rewarded with water dispensed from the central lick port. (D, E) Illustration of the optogenetic suppression design, in VGAT-ChR2 transgenic mice. Optogenetic suppression (blue light-on) of wMC or S1 was performed on 33% of trials, randomly interleaved with control (light-off) trials. (F) Features of optogenetic stimulation: the optogenetic light (blue triangle) starts ramping up before the whisker stimulus, peaks after stimulus onset, and ramps down during the response window. A 10 Hz pulsing visual mask, present on all trials, fully covers the duration of optogenetic suppression. Modulation index is used to quantify changes in response rates, with positive modulation indices indicated higher response rates during optogenetic suppression. (G) Hit rates and false alarm rates for optogenetic light-on trials and control light-off trials across all sessions for suppression of target-aligned wMC (1^st^ column), distractor-aligned wMC (2^nd^ column), target-aligned S1 (3^rd^ column), and distractor-aligned S1 (4^th^ column). The top row indicates the site of optogenetic suppression. The middle row displays hit rates (magenta) and false alarm rates (orange), each circle reflecting one behavioral session. The bottom row displays the modulation indices calculated based on the middle row data. Throughout the article, p values are indicated as: n.s. (p>=0.05), *(0.01<=p<0.05), **(0.001<=p<0.01), or ***(p<0.001).

### Either target-aligned or distractor-aligned wMC suppression increases false alarms

We tested the impacts of wMC optogenetic suppression on task performance. Focal suppression was achieved by optical activation of GABAergic interneurons in either target-aligned (contralateral to target whisker stimuli) or distractor-aligned (contralateral to distractor whisker stimuli) wMC (Fig. 1 D,F). Suppression was initiated 100-200 ms before stimulus onset, and was robust and stable throughout the post-stimulus lockout window (Fig. 3A,B, Fig. 4A). Control trials (light-off) and wMC suppression trials (light-on) were randomly interleaved. Interestingly, suppression of target-aligned or distractor-aligned wMC increased large amplitude (Fig. 1G) and small amplitude (Supplementary Fig. 1A,B) stimulus false alarm rates (target-aligned: large stimulus false alarm rate: n=28 sessions, light-off: 0.33±0.02, light-on: 0.54±0.04, paired test: p=2.70×10^-7^, MI=0.26±0.04, one sample t test: p=6.00×10^-7^; distractor-aligned: large stimulus false alarm rate: n=19 sessions, light-off: 0.26±0.03, light-on: 0.46±0.05, paired test: p=5.00×10^-4^, MI=0.27±0.06, one sample t test: p=2.75×10^-4^). Across the same sessions we did not observe significant changes in large stimulus hit rates (target-aligned: large stimulus hit rate: n=25 sessions, light-off: 0.95±0.03, light-on: 0.96±0.02, paired test: p=0.67, MI=0.01±0.01, one sample t test: p=0.434, distractor-aligned: large stimulus hit rate, n=15 sessions, light-off: 0.93±0.02, light-on: 0.93±0.02, paired test: p=0.92, MI=0.00±0.01, one sample t test: p=0.8876) (Fig. 1G) or spontaneous response rates (Supplementary Fig. 1). Suppression of target-aligned or distractor-aligned wMC caused trends towards increased small amplitude hit rates (with statistical significance depending on calculation method, Supplementary Fig. 1A,B). These data support wMC function in suppressing behavioral responses, most robustly for distractor stimuli.

**Fig. 2:**
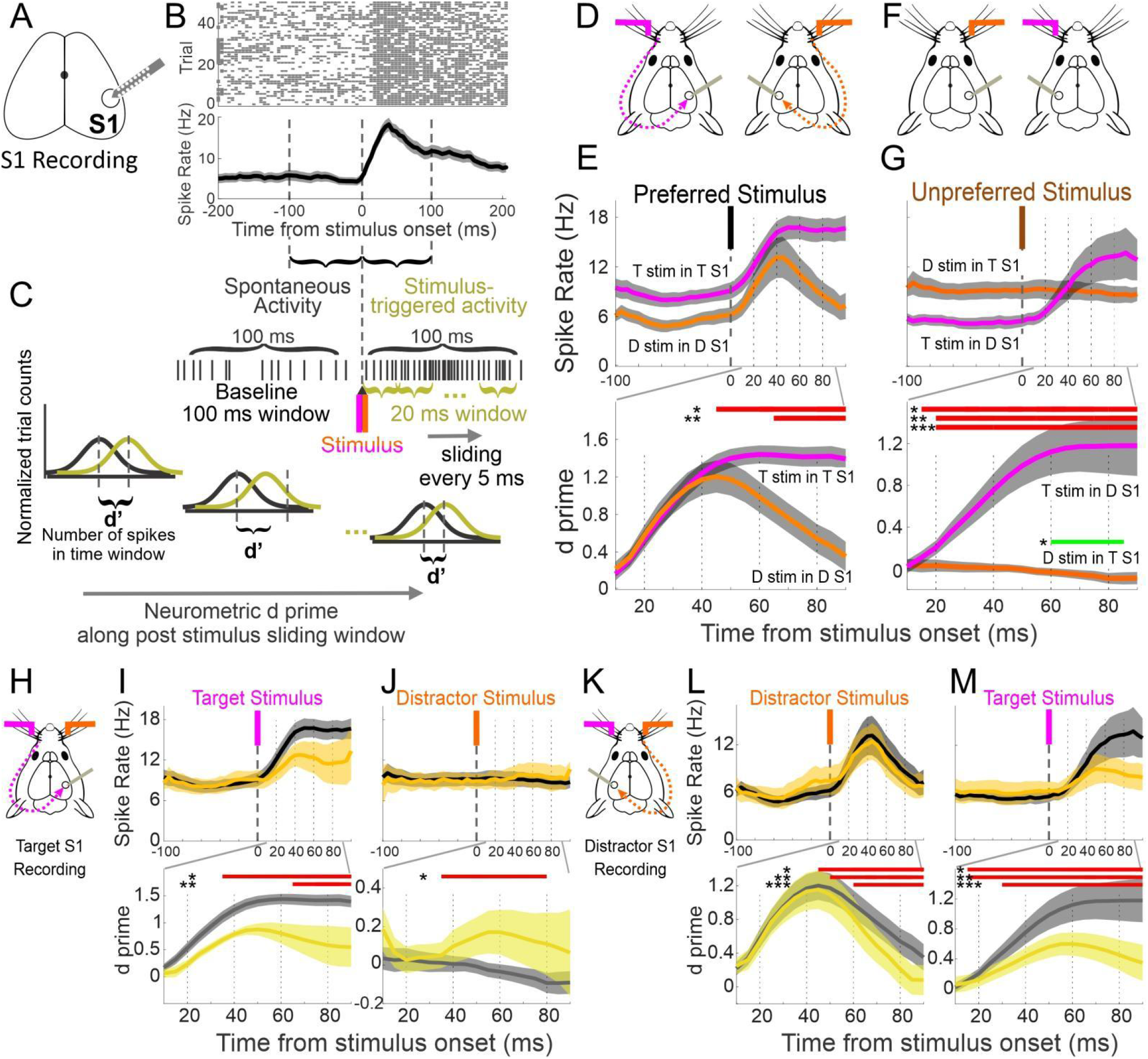
Distractor stimulus encoding is suppressed in target-aligned S1 during task engagement. (A) Illustration of extracellular silicon probe recording of S1. (B) Example session of S1 multi-unit spiking activity in response to preferred whisker stimuli, from one awake behaving recording session. Top plot: raster plots of neuronal spiking for multiple trials. Bottom plot: averaged neuronal spike rates from the data above. (C) Illustration of our method of calculating neuronal d prime. Neuronal activity from 100ms windows before and after whisker stimulus onset are assigned as spontaneous activity (stimulus absent) and stimulus-triggered activity (stimulus present), respectively. To generate a neurometric time series, we use a 20 ms sliding window across the 100ms post-stimulus period (with duration-matched samples from the pre-stimulus window, see Methods). (D-G) Analyses of recordings in expert mice during task performance. (D) Experimental design illustration for [E]: recording in target-aligned or distractor-aligned S1 to their preferred stimuli. (E) Neuronal activity (top) and d prime (bottom), overlaying data from target stimulus responses in target-aligned S1 (magenta) and distractor stimulus responses in distractor-aligned S1(orange). The d prime time series (bottom) only reflects the post-stimulus window. Data are grand averages, combined from all recording sessions (n=17 target-aligned recording sessions, n=10 distractor-aligned recording sessions). The red bars indicate the epochs in which d prime from two recording groups are significantly different from each other; same indication for [G,I,J,L,M]. (F) Experimental design illustration for [G]: recording in target-aligned or distractor-aligned S1 to their unpreferred stimuli. (G) As same as [E] but distractor stimulus responses in target-aligned S1 (orange) and target stimulus responses in distractor-aligned S1(magenta). The green bar indicates the epochs in which d prime for distractor stimulus encoding in target-aligned S1 is significantly below zero. (H-M) Comparisons of S1 neuronal activity between engaged and disengaged recording sessions. (H) Experimental design illustration for [I, J]. (I) Neuronal activity (top) and d prime (bottom) in target-aligned S1 for preferred stimuli (task-engaged, black; task-disengaged, yellow) (J) As same as [I] but for unpreferred stimuli. (K-M) As same as [H-J] but for distractor-aligned S1 recordings.

**Fig. 3:**
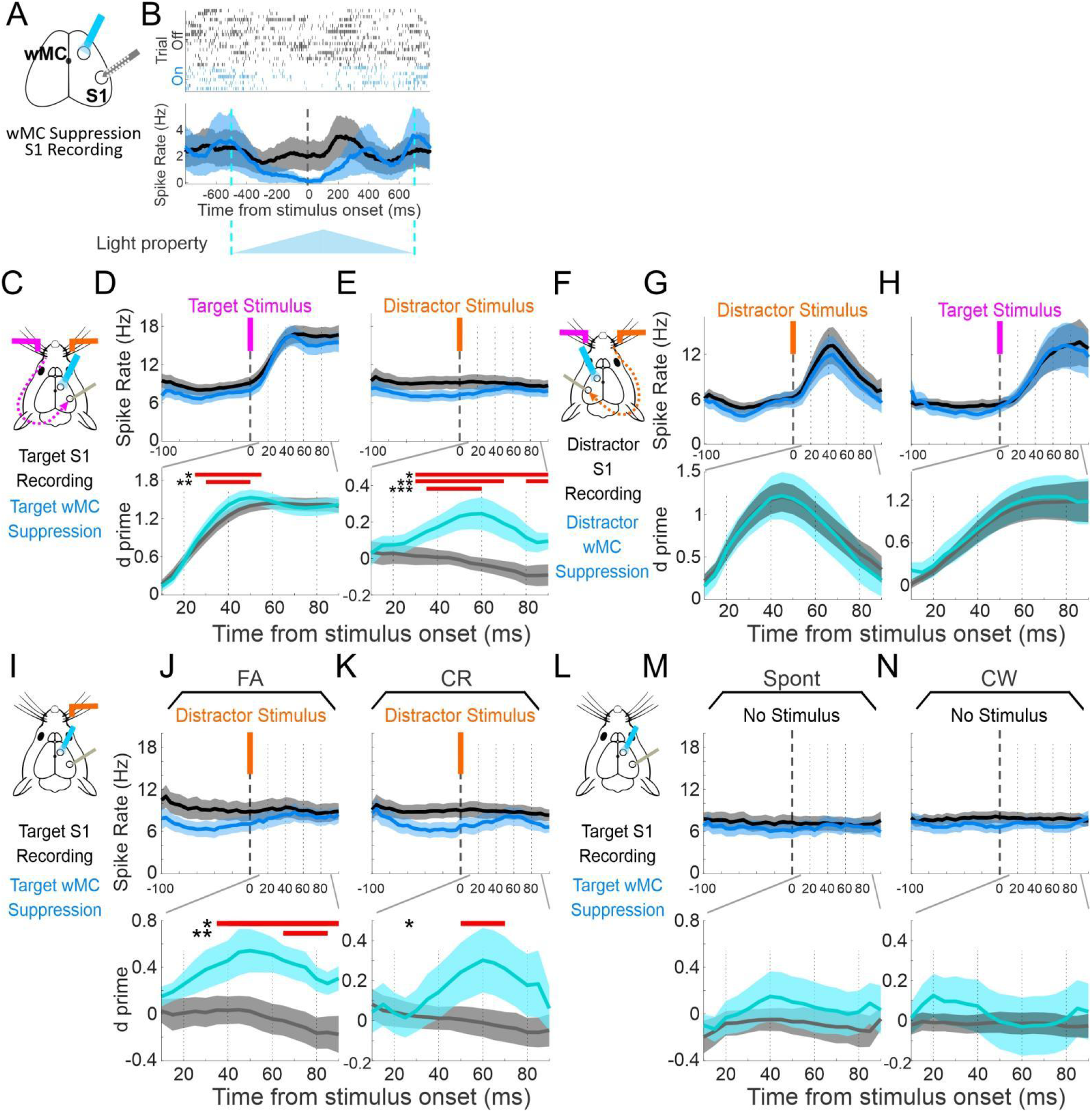
Suppressing target-aligned wMC increases distractor stimulus encoding in target-aligned S1. (A) Illustration of simultaneous optogenetic suppression of wMC and extracellular silicon probe recording of S1. (B) Example effects of wMC suppression on S1 multi-unit spiking activity, from one anesthetized recording session (only catch trials are shown). Top plot: raster plots of neuronal spiking for multiple trials. Middle plot: averaged neuronal spike rates from the data above. Black, light-off control trials; dark blue, light-on optogenetic suppression trials. Bottom plot: time course of optogenetic stimulation. (C-H) Analyses of recordings in expert mice during task performance. (C) Experimental design illustration for [D, E]: recording in target-aligned S1, suppression of target-aligned wMC, target stimulus is preferred and distractor stimulus is unpreferred. (D, E) Neuronal activity (top) and d prime (bottom) of target-aligned S1 neurons during control trials (light-off, black/gray) and wMC suppression trials (light-on, dark blue/light blue) in response to target [D] or distractor [E] stimuli. Data are grand averages, combined from all recording sessions. The red bars indicate the epochs in which d prime during wMC suppression is significantly different from control trials. (F) Experimental design illustration for [G, H]: recording in distractor-aligned S1, suppression of distractor-aligned wMC, distractor stimulus is preferred and target stimulus is unpreferred. (G, H) As same as [D, E] but for distractor-aligned S1 neurons during distractor-aligned wMC suppression. Note, the most robust effect of wMC suppression is increased distractor stimulus encoding in target-aligned S1 [E]. For neuronal activity perturbations across all stimulus categories, see Supplementary Fig. 3. (I-N) Behavioral response or wMC suppression alone does not account for increased distractor-evoked responses in target-aligned S1. Data are presented as above. (I) Diagram of target-aligned S1 recordings and target-aligned wMC suppression for distractor trials. Comparison of wMC suppression and control conditions on false alarm (FA) trials (J) and correct rejection (CR) trials (K). (L-N) Same as [I-K] but for catch trials without whisker stimuli, showing spontaneous responding (Spont) and correct withhold (CW) trials.

**Fig. 4:**
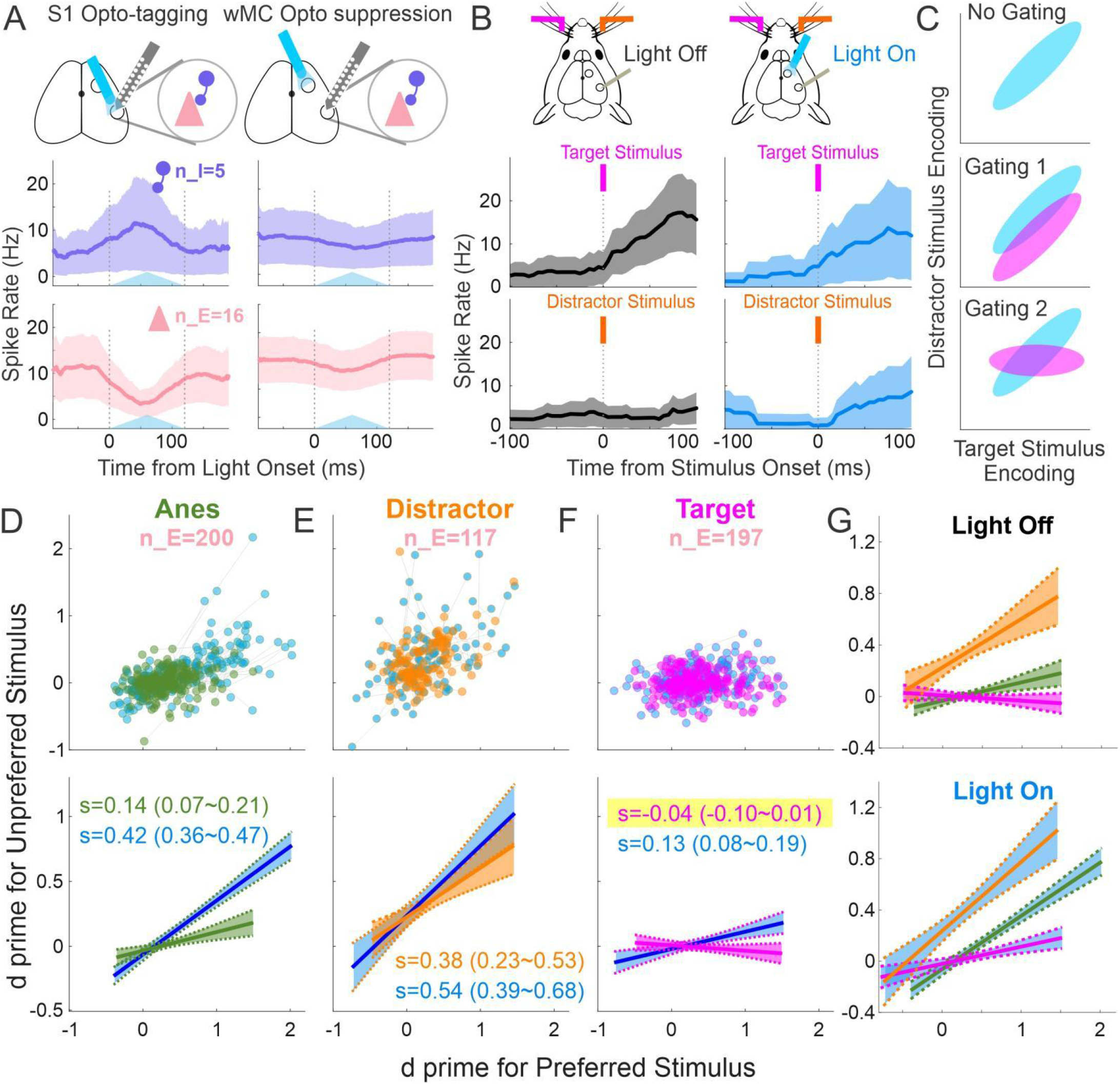
Asymmetric stimulus selectivity in target-aligned and distractor-aligned S1 and its modulation by wMC. (A) Example responses of S1 putative excitatory and inhibitory neurons to S1 opto-tagging (left) and wMC optogenetic suppression (right). Top row: for S1 opto-tagging, a light fiber is positioned above the S1 recording probe. For wMC suppression, a second light fiber is positioned above wMC. Middle and bottom plots: average spiking activity recorded from one example awake behaving session. Purple traces indicate activity of putative inhibitory neurons during S1 opto-tagging (left) and the same neurons during wMC opto suppression (right). Pink traces indicate activity of putative excitatory neurons during S1 opto-tagging (left) and the same neurons during wMC opto suppression (right). The blue triangles reflect the time course of optical stimulation. (B) Target stimulus and distractor stimulus evoked responses of an example putative excitatory neuron in target-aligned S1. Left: during light-off control trials, this neuron displayed a large stimulus-triggered response after target stimuli (top) but no response after distractor stimuli (bottom). Right: in light-on wMC suppression trials, the same neuron displayed robust stimulus-triggered responses for both target (top) and distractor (bottom) stimuli. (C) Illustration of potential mechanism of distractor stimulus gating plotted in stimulus selectivity space, with target stimulus encoding on the x-axis and distractor stimulus encoding on the y-axis. Blue distributions reflect a population of target-aligned S1 neurons without distractor gating. Magenta distributions reflect the same population of neurons with distractor gating. Top: without gating, neurons in S1 robustly encode both target and distractor stimuli. Middle: one potential mechanism of gating is for the population to display the same stimulus selectivity correlation, yet with reduced distractor stimulus encoding throughout the population. Bottom: a second potential mechanism of gating, with a rotation in the selectivity space resulting in a decorrelation of target and distractor stimulus encoding throughout the population. (D-F) Putative excitatory neurons recorded (D) under anesthesia, (E) in distractor-aligned S1 in awake behaving mice, and (F) in target-aligned S1 in awake behaving mice. In each plot, the top scatter plot shows single neurons’ d prime for the preferred stimulus (contralateral to the recording site) vs the unpreferred stimulus (ipsilateral to the recording site). The green, orange, and magenta dots indicate the values with wMC intact (control, light-off) and the blue dots indicate the values of the same neurons with wMC suppression (light-on). Lines connect the same neurons under light-off and light-on conditions. The bottom plots show the linear fits of the single unit data above. The numbers indicate the slopes (s) of the fitted lines [fit (95% confidence interval)]. (G) The overlay of the light-off control (top) and light-on wMC suppression (bottom) linear fits, re-plotted from the second row of [D-F].

To determine the specificity of this effect, we additionally tested the impacts of optogenetic suppression of S1 (instead of wMC) on task performance (Fig. 1E). Suppression of target-aligned S1 resulted in significantly decreased small amplitude hit rates (Supplementary Fig. 1C) and trends towards decreased hit rates and false alarm rates for large amplitude stimuli (Fig. 1G). Decreased response rates during S1 suppression contrast with increased response rates during wMC suppression (comparing suppression of target-aligned wMC and target-aligned S1, two sample t test: large stimulus hit rate, p=0.02; large stimulus false alarm rate, p=5.67×10^-4^). These data suggest different functions of wMC and S1 in this task, with wMC suppressing and S1 promoting behavioral responses.

### Suppression of distractor stimulus propagation during task engagement

We recorded spiking activity of single units in layer 5 S1 during task performance (Fig. 2A,B). First, we analyzed population (summation of single units recording simultaneously in each session) stimulus encoding using neurometric analyses (Fig. 2C). We refer to preferred stimuli (target stimuli for target-aligned S1, distractor stimuli for distractor-aligned S1) and unpreferred stimuli (distractor stimuli for target-aligned S1, target stimuli for distractor-aligned S1). In awake behaving mice during task engagement, target-aligned and distractor-aligned S1 encoded their preferred stimuli robustly and at short latency, peaking at 40∼45 ms post-stimulus (Fig. 2E). As shown in the neurometric (d prime) plots (Fig. 2E, bottom), these signals overlap until approximately 40 ms, after which the target stimulus encoding remains elevated and the distractor stimulus encoding decreases back to baseline.

Distractor-aligned S1 also showed robust encoding of unpreferred (target) stimuli (Fig 2G, magenta traces), peaking at approximately 60∼80 ms post-stimulus. This longer latency is consistent with a longer pathway for unpreferred (ipsilateral) stimuli to reach S1. In marked contrast, we observed a slight negative encoding of distractor stimuli in target-aligned S1 (Fig 2G, orange traces) (−0.02±0.02, averaged over the 100 ms window, n=17 sessions; one sample t test for all 17 sessions in each 20 ms sliding window, 60-80ms: p=0.05; 65-85ms: p=0.04). Accordingly, unpreferred stimulus encoding in distractor-aligned S1 was significantly larger than in target-aligned S1 (neurometric averaged across 100 ms, two sample t test, p=9.54×10^-7^), with this difference emerging immediately post-stimulus. Thus, ignoring distractor stimuli is correlated with suppressed encoding of distractor stimuli in target-aligned S1, compared to the robust encoding of target stimuli in distractor-aligned S1.

Next, we determined which aspects of preferred and unpreferred stimulus encoding were dependent on task engagement. Neuronal activity during task disengagement were obtained from the same expert mice, after they stopped responding within a session (presumably due to satiety). Generally, disengagement led to reduced stimulus encoding (Fig 2H-M). This reduction was observed for target stimuli in target-aligned S1 (Fig 2I), distractor stimuli in distractor aligned S1 (Fig 2L), and target stimuli in distractor-aligned S1 (Fig. 2M). In marked contrast, disengagement led to an increase in distractor encoding in target-aligned S1 (Fig 2J). In summary, we observed a strong asymmetry in the propagation of target and distractor stimuli across hemispheres with a suppression of distractor stimulus response propagation into target-aligned S1 (Fig 2G), which was dependent on task engagement (Fig 2J).

To verify this observation using a different recording method, we analyzed widefield Ca^2+^ - sensor imaging data from different mice performing the same behavioral task (Aruljothi et al., 2020). We observed a similar asymmetric interhemispheric propagation in our imaging data (Supplementary Fig 2). We found that target stimuli induced significant activation of distractor-aligned S1 (n=40 sessions, d prime 0.14±0.03, one sample t-test p=0.0002). In contrast, distractor stimuli induced significant suppression of target-aligned S1 (n=40 sessions, d prime −0.07±0.02, one sample t-test p=0.007; paired t-test comparing target and distractor propagation, p=1.8×10^-5^).

### Suppressing target-aligned wMC increases distractor stimulus encoding in target-aligned S1

One of the main output projections from wMC is to S1, through both the direct cortical feedback pathway and indirect cortico-thalamo-cortical pathways (Aronoff et al., 2010; Manita et al., 2015; Mao et al., 2011; Sato & Svoboda, 2010; White & DeAmicis, 1977). Therefore, we sought to determine the consequences of wMC suppression on sensory encoding in S1. While we recorded S1 neuronal activity described above, we also applied interleaved optogenetic wMC suppression (Fig. 3A). wMC suppression reduced baseline firing rates for all conditions (Fig. 3B, characterized further below and in Fig. 5). Here we focus on the effects of wMC suppression on stimulus encoding, as determined by population neurometric analyses. Specifically, we conducted these analyses to assess how target-aligned wMC suppression changes target-aligned S1 encoding of target and distractor stimuli (Fig. 3C-E), and how distractor-aligned wMC suppression changes distractor-aligned S1 encoding of distractor and target stimuli (Fig. 3F-H).

**Fig. 5:**
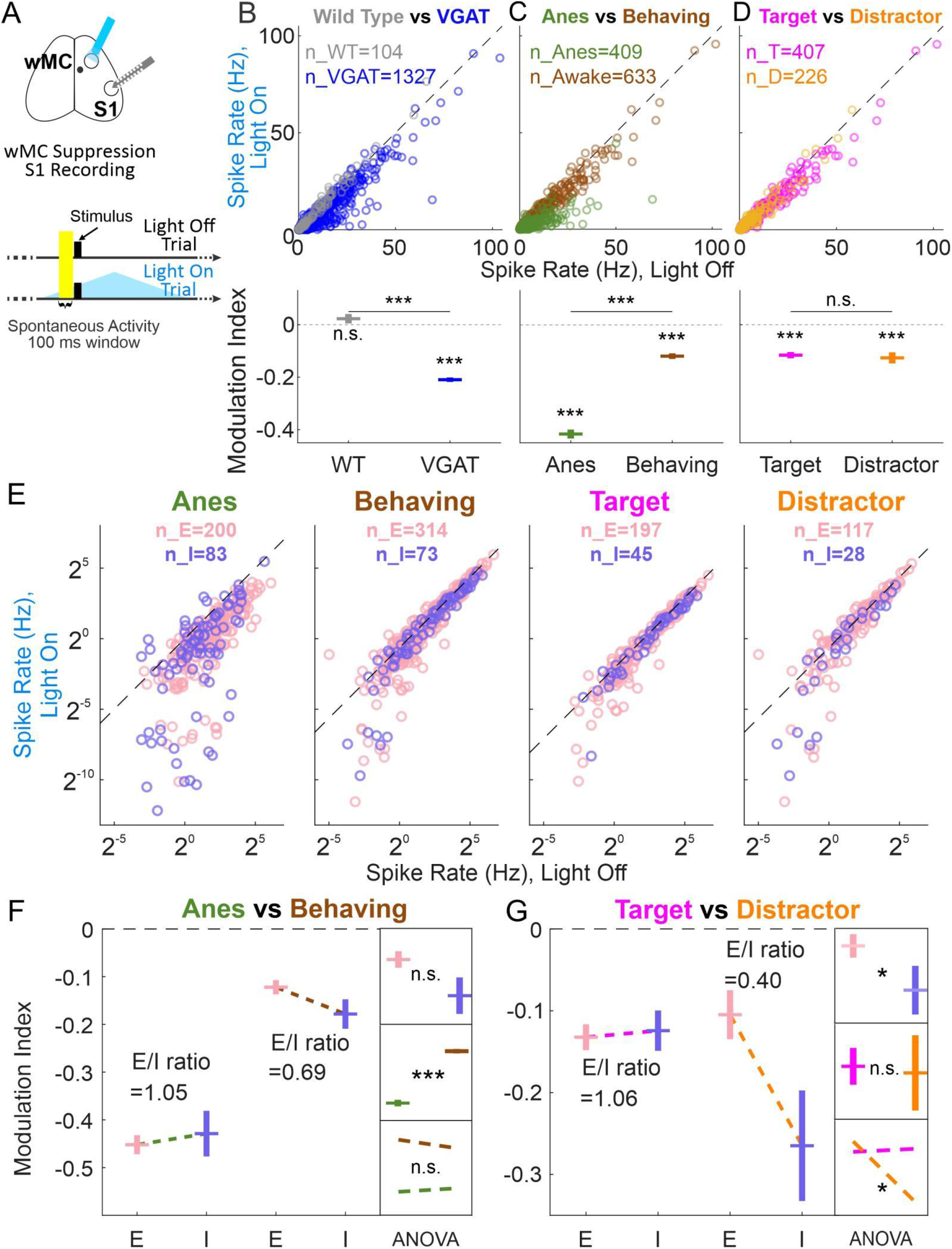
wMC proactively drives S1 excitatory and inhibitory neurons in a context-dependent manner. (A) Experimental design, as described previously (S1 opto-tagging was also performed but not shown here). Neuronal activity within the 100 ms pre-stimulus window (highlighted) was used to determine pre-stimulus wMC modulation. (B) Top: S1 pre-stimulus spike rates from wild type (gray) and VGAT-ChR2 (blue) mice. Each circle indicates one neuron. Bottom: modulation index calculated based on the top plot. Above each data point are results from one-sample t tests to determine differences from zero within each population, and lines above the data points are results from two-sample t tests to determine differences between populations. (C) As same as [B] but for anesthetized (green) and awake behaving sessions (brown, including both target-aligned and distractor-aligned S1) of VGAT-ChR2 mice. (D) As same as [B] but for awake behaving sessions of VGAT-ChR2 mice in target-aligned S1 (magenta) and distractor-aligned S1 (orange). (E) As same as [B-D] but separated for excitatory (pink) and inhibitory (purple) neurons under different contexts. (F) Statistical analyses for anesthetized vs awake behaving data. Big box on the left: modulation indices based on the spike rates in [E]. Small boxes on the right: results of two-way ANOVA for the modulation indices of anesthesia and awake behaving contexts and excitatory and inhibitory neuron types: top: no significant main effect of neuron type; middle: a significant main effect for context; bottom: no significant interaction effect. (G) Statistical analyses for target-aligned S1 vs distractor-aligned S1 awake behaving data, organized as in [F]. ANOVA results: top: a significant main effect for neuron type; middle: no significant main effect of context (alignment); bottom: a significant interaction effect.

Using the same method described above for the task engaged vs disengaged comparison, here we report the effects of wMC suppression on S1 stimulus encoding by comparing optogenetic light-off (black and gray) and light-on (dark blue and light blue) conditions (Fig. 3C-H). wMC suppression transiently increased target stimulus encoding in target-aligned S1 (Fig. 3D), which we interpret as wMC marginally suppressing target stimulus encoding under control conditions. wMC suppression did not significantly impact distractor-aligned S1 encoding of distractor (preferred) (Fig. 3G) or target (unpreferred) stimuli (Fig. 3H). However, wMC suppression did significantly increase target-aligned S1 encoding of distractor (unpreferred) stimuli (Fig. 3E) (two sample t test, p<0.05 during 30-100 ms post stimulus window). This effect was present from 30-100 ms post-stimulus, well before the response time (minimum 200 ms), and similar to the latency of unpreferred stimulus encoding in distractor-aligned S1. Importantly, this increase in target-aligned S1 encoding of distractor stimuli was present regardless of trial outcome (false alarm or correction rejection, Fig. 3I-K) and was not present with optical stimulation alone (i.e., on catch trials) (Fig. 3L-N). From these data, we conclude that the main impact of wMC on sensory encoding is to suppress the propagation of distractor stimulus responses into target-aligned S1. Moreover, this neurometric finding is consistent with, and indeed may contribute to, the increased false alarm rates during wMC suppression (Fig. 1G).

### Target-aligned wMC decorrelates target and distractor stimulus encoding in target-aligned S1 excitatory neurons

In the analyses described above we report population activity (from summed single units). Here, we report single unit analyses, segregated into putative excitatory and inhibitory neurons based on opto-tagging (Fig. 4A and Supplementary Fig. 5). Below, we focus our analyses on putative excitatory neurons, as the output of S1.

First, we present the activity of one example excitatory neuron in target-aligned S1 (Fig. 4B). In control (light-off) conditions (Fig. 4B, left), this neuron responds robustly to target stimuli, but not to distractor stimuli. During wMC suppression (Fig. 4B, right), responses to target stimuli are maintained, yet now we observe increased responses to distractor stimuli. We interpret this finding as reduced stimulus selectivity, in which this neuron increases its apparent receptive field size to now include distractor (unpreferred) inputs. This suggests that an important function of wMC is to gate out distractor stimulus responses from target-aligned S1 neurons.

We recognize two potential mechanisms by which wMC may gate distractor stimulus responses in target-aligned S1 (Fig. 4C). First, there could be a shift of the entire population, such that throughout the distribution of S1 neurons there is a proportional reduction of distractor encoding (gating model 1, Fig. 4C, middle). Second, there could be a rotation in the distribution of S1 neurons, such that encoding of target and distractor stimuli are decorrelated in individual neurons (gating model 2, Fig. 4C, bottom). To distinguish between these mechanisms, we plotted the distribution of preferred (contralateral) and unpreferred (ipsilateral) stimulus encoding (neuronal d prime over a 0-100ms post stimulus window compared to pre-stimulus, see Methods) from target-aligned S1 neurons, distractor-aligned S1 neurons, and from S1 neurons in naïve, anesthetized mice (Fig. 4D-G). We first describe stimulus selectivity profiles in control light-off conditions. In anesthetized mice (Fig. 4D,G, green), S1 neurons displayed a slight positive correlation, such that the neurons with the strongest encoding of preferred (contralateral) stimuli also positively encoded unpreferred (ipsilateral) stimuli (95% confidence interval of the slope of the linear fit: 0.07-0.21, for n=200 single units). During task performance, distractor-aligned and target-aligned S1 neurons displayed remarkably different stimulus selectivity profiles. Distractor-aligned S1 neurons were less selective than the anesthetized population, with a more positive slope to the linear fit of their stimulus response correlations (95% confidence interval of the slope of the linear fit: 0.23-0.57, for n=117 single units) (Fig. 4E,G, orange). In contrast, target-aligned S1 neurons were more selective than the anesthetized population, with a flat, trending to negative, slope to the linear fit of their stimulus response correlations (95% confidence interval of the slope of the linear fit: −0.10-0.01, for n=197 single units) (Fig. 4F,G, magenta). That is, across target-aligned S1 neurons, target and distractor stimulus encoding is decorrelated.

Next, we tested the impact of wMC suppression on stimulus selectivity, starting with target-aligned S1. As shown in Fig. 4F (blue), wMC suppression caused a rotation in the stimulus selectivity of target-aligned S1 neurons, such that now there was a positive slope to the population stimulus response correlation (95% confidence interval of the slope of the linear fit: 0.08-0.19). Thus, wMC contributes to distractor response suppression in target-aligned S1 by modulating single unit sensory selectivity. This rotation in stimulus correlation space (Fig. 4F bottom) is consistent with the ‘gating 2’ model shown in Fig. 4C, as opposed to a shift in distractor stimulus encoding across the population (‘gating 1’ model).

wMC suppression also caused a positive rotation in the stimulus selectivity space for the other conditions: significant for anesthetized recordings (Fig. 4D), and a non-significant trend for distractor-aligned S1 (Fig. 4E). These data may suggest that suppression of unpreferred stimuli is a general, non-context dependent, function of wMC. However, further analyses indicate that stimulus selectivity and its modulation by wMC is indeed task modulated and context dependent. First, the effects of wMC suppression were different for putative inhibitory neurons (Supplementary Fig. 6A,B). In anesthetized mice, there was a trend for wMC suppression to increase the slope of stimulus correlation (reflecting reduced selectivity) of putative inhibitory neurons. Yet, there was a trend in the opposite direction for putative inhibitory neurons in target-aligned and distractor-aligned S1 during task performance (reduced slope, increased stimulus selectivity) (Supplementary Fig. 6A). Moreover, when mice were disengaged from the task, differences in stimulus selectivity were less pronounced between target-aligned and distractor-aligned S1 (Supplementary Fig. 6C-F). Importantly, when mice were disengaged, wMC suppression in target-aligned S1 did not change stimulus selectivity (Supplementary Fig. 6C). Altogether, these data suggest that asymmetric modulation of stimulus selectivity in S1 is a robust feature of selective detection, and that wMC contributes to this modulation.

The wMC-mediated decorrelation of target and distractor encoding in target-aligned S1 excitatory neurons is likely to be critical for distractor response suppression in this task. During wMC suppression, due to the positive correlation across single units for sensory encoding of target and distractor stimuli, it would be difficult for a down-stream reader to distinguish between a small target stimulus and a large distractor stimulus, and thus could account for the increased false alarm rates. We tested this hypothesis using a linear classier to mimic the downstream reader. First, we chose the half of the target-aligned S1 excitatory neurons with the largest target stimulus encoding, as the putative population signaling target detection to the downstream region. Next, we trained a classifier on single trial data from this ensemble, to distinguish between target and distractor trials. We then tested the classifier on both hold-out control trials and wMC suppression trials, from the same neuronal population (see methods). We found that this simulated downstream reader was nearly perfect at correctly classifying control data as target and distractor trials (error rate, 0.55%). In marked contrast, the downstream reader error rate was substantial for wMC suppression data (20.31%, Supplementary Fig. 7). Notably, the error types for wMC suppression data partially matched the behavioral data, demonstrating higher ‘false alarm rates’ (mis-classification of distractor as target trials) than ‘miss rates’ (mis-classification of target as distractor trials) (30.09% vs 10.52%, Supplementary Fig. 7).

### wMC proactively modulates S1 activity in a context-dependent manner

Top-down modulations of sensory cortex can occur in two fundamentally different ways. Top-down modulations may be stimulus-evoked and reactively exert a functional feedback signal (Angelucci et al., 2002; Hupé et al., 1998; Manita et al., 2015; Moore & Armstrong, 2003; Nassi et al., 2013). Alternatively, top-down modulations may be in place before a stimulus arrives, proactively setting the initial condition for goal-directed sensory processing (Zagha, 2020). These last sets of analyses were to determine whether there is evidence for proactive (pre-stimulus) wMC modulation of S1 spike rates (SR). For these analyses, we considered the last 100 ms pre-stimulus, in which we could achieve a relatively stable baseline during wMC optogenetic suppression (Fig. 3B, Fig. 5A). We calculated the pre-stimulus modulation index for each neuron as MI= (light_on_SR – light_off_SR) / (light_on_SR + light_off_SR). As such, MI < 0 indicates that wMC suppression reduces spiking (alternatively, that wMC normally drives spiking) in that neuron.

For recording conditions of anesthetized, awake behaving target-aligned, and awake behaving distractor-aligned, wMC robustly drives pre-stimulus spiking in S1 neurons (Fig. 5B-D). This effect was not present in wild type control mice lacking channelrhodopsin expression, and therefore is unlikely to be due to artifacts of optogenetic stimulation (modulation index (MI) of each group, one sample t test p values of each group, and two sample t test p values between groups: wild-type: 0.02±0.02, p=0.21; VGAT: −0.21±0.01, p=7.83×10^-112^; WT vs VGAT: 5.97×10^-14^) (Fig. 5B). Interestingly, wMC modulation of S1 pre-stimulus activity under anesthesia was larger than during task performance (MI of each group, one sample t test p values of each group, and two sample t test p values between groups: anesthesia: −0.42±0.02, p=4.87×10^-82^; awake behaving: - 0.12±0.01, p=1.44×10^-26^; anesthesia vs awake behaving: p=4.72×10^-49^) (Fig. 5C). The modulation indices were similar in awake behaving target-aligned and distractor-aligned S1 (MI of each group, one sample t test p values of each group, and two sample t test p values between groups: target: - 0.12±0.01, p=2.03×10^-20^; distractor: −0.13±0.02, p=8.47×10^-9^; target vs distractor: 0.6457) (Fig. 5D), indicating similar levels of overall wMC drive.

To gain further insights into changes in local circuits, we analyzed wMC modulation separately for putative excitatory and inhibitory S1 neurons. Both excitatory and inhibitory S1 neurons were significantly driven by wMC in anesthetized and awake behaving mice, both target-aligned and distractor-aligned (paired t test: anesthetized excitatory: p= 8.60×10^-17^; anesthetized inhibitory: p= 8.91×10^-8^; awake excitatory: p=3.10×10^-12^; awake inhibitory: p=9.70×10^-6^; target-aligned excitatory: p=1.19×10^-11^; target-aligned inhibitory: p=2.26×10^-4^; distractor-aligned excitatory: p=0.02; distractor-aligned inhibitory: p=0.01) (Fig. 5 E-G). For each condition, we calculated the E/I modulation ratio, as the MI of excitatory neurons divided by the MI of inhibitory neurons. E/I ratio for anesthetized and target-aligned conditions were remarkably balanced, yielding ratios near 1. In contrast, the distractor-aligned E/I ratio was 0.40, reflecting a preferential drive of wMC onto putative inhibitory neurons (Fig. 5G). Two-way ANOVA analyses supported these conclusions, yielding a significant main effect for behavioral context comparing anesthetized and awake behaving conditions (anesthetized vs awake, p=1.23×10^-24^) (Fig. 5F). Comparing target-aligned and distractor-aligned conditions, we found a main effect for cell-type (p=0.03) and an interaction effect between alignment and cell-type (p=0.02) (Fig. 5G). These data indicate that wMC modulation of S1 is proactive (influencing pre-stimulus activity levels) and context-dependent (for anesthesia vs awake behaving and target vs distractor alignment).

## Discussion

The major significance of this study is identifying a mechanism by which frontal cortex suppresses behavioral and neuronal responses to distractor stimuli. Within frontal cortex, we focused on a region that is part of the sensorimotor cortical whisker system, termed ‘whisker motor cortex’ or wMC. Using optogenetic suppression, we demonstrated that wMC robustly contributes to distractor response suppression (Fig. 1G, Supplementary Fig. 1A,B). From S1 single unit recordings in behaving mice we revealed an asymmetry in stimulus selectivity, in which distractor-aligned S1 neurons respond to both distractor and target stimuli while target-aligned S1 neurons respond selectively to target stimuli (Fig. 2D-G, Fig. 4G). By combining wMC optogenetic suppression with S1 single unit recordings in behaving mice we demonstrated that wMC contributes to target-aligned S1 stimulus selectivity, such that with wMC suppression S1 neurons more strongly encode distractor stimuli (Fig. 3E, Fig. 4F). This reduction in distractor gating likely underlies, at least in part, the increase in false alarm rate during wMC suppression (discussed further below). Lastly, we provide strong evidence that wMC modulation of S1 is context-dependent and proactive (present prior to stimulus onset). Our study establishes a tractable and behaviorally-relevant framework for how top-down modulations of sensory propagation implement sensory gating to mediate distractor response suppression.

For combined behavioral-physiological studies, an important consideration is whether the task-related neuronal signals could be fully or partially accounted for by movement (Zagha et al., 2022). A specific concern for our study is whether the apparent asymmetric stimulus selectivity, and its modulations by wMC, is instead reflecting movement on ‘Go’ trials in our Go/No-Go task design. We believe this is unlikely for four reasons. First, differences in selectivity (between target-aligned S1 and distractor-aligned S1, between control trials and wMC suppression trials) are observable by 30 ms post-stimulus onset. This is well before the response time (>200 ms), before the onset of uninstructed whisker movements that are part of the ‘Go’ motor sequence (>80 ms, Aruljothi et al., 2020, and data not shown), and well before the emergence of significant choice probability in S1, which includes the contributions of ‘Go’ movements (>165 ms, Zareian et al., 2021). Second, increases in distractor responses in target-aligned S1 are present irrespective of trial outcome (false alarm or correct rejection), and are not present on catch trials (i.e., in the absence of distractor stimuli) (Fig. 3I-N). Third, target-aligned S1 neurons during task performance are more sensory selective than S1 neurons under anesthesia, during which movements are largely suppressed. Fourth, wMC suppression causes a rotation in stimulus selectivity (Fig. 4F) rather than a shift of the population, as would be expected from increases in arousal or movement (Carsen et al., 2019). For these reasons we believe that our reported modulations of stimulus selectivity are truly reflecting sensory processing rather than movement confounds.

A major framework of top-down cortical modulation is enhancing sensory responses of neurons aligned to the top-down inputs (center) and suppressing sensory responses of neurons unaligned (surround), as measured for the optimal (preferred) stimulus for each population (Angelucci et al., 2002; Hupé et al., 1998; Keller et al., 2020; Moore & Armstrong, 2003; Zhang et al., 2014). Data from our study, however, do not support this framework. Notably, wMC suppression had only minor impacts on encoding of preferred stimuli (Fig. 3D,G). This was observed for both large and small amplitude stimuli (Supplementary Fig. 3), and therefore is unlikely due to response saturation. Our data are more consistent with top-down cortical modulation suppressing sensory processing (Fritz et al., 2003; Fritz et al., 2010; Moran & Desimone, 1985; Nassi et al., 2013), yet specifically for the propagation of distractor stimulus responses into target-aligned cortex.

We speculate as to the cellular and circuit mechanisms underlying these sensory response modulations and their impacts on behavioral responses. Our analyses of pre-stimulus spike rates were notable for a preferential wMC activation of distractor-aligned S1 inhibitory neurons (Fig. 5G). While this increased inhibitory drive did not substantially suppress distractor stimulus encoding in distractor-aligned S1, we hypothesize that these inhibitory neurons may prevent the spread of distractor evoked responses into target-aligned cortex. Additionally, we have considered why the propagation of distractor responses into target-aligned S1 would impair distractor response suppression. In this selective detection task, we suspect that behavioral responses are conditioned on the activation of target-aligned regions, including from target-aligned S1 (Supplementary Fig. 1C). However, in naïve rodents, whisker-evoked responses propagate widely and across hemispheres (Aronoff et al., 2010; Aruljothi et al., 2020; Ferezou et al., 2007; Pala & Stanley, 2022; Reig and Silberberg, 2016). We reason that it would be difficult for a downstream reader to distinguish between target-aligned S1 activations that propagated bottom-up from target stimuli vs target-aligned S1 activations that propagated laterally from distractor stimuli. Thus, an important function of top-down signals would be to restrict the propagation of distractor stimuli into target-aligned S1, thereby increasing the likelihood (posterior probability) that target-aligned S1 activations reflect the occurrence of a target stimulus. Without top-down restrictions of distractor propagation, downstream readers would be more likely to respond incorrectly to target-aligned S1 activations evoked by distractor stimuli (as simulated in Supplementary Fig. 7).

We note that wMC modulation of S1 is only a partial component of both wMC function and S1 dynamics. wMC suppression not only increased false alarm rates, but also substantially increased premature responses (during the 200 ms post-stimulus lockout window) (Supplementary Fig. 1A,B). We do not find strong evidence for increased premature responses being accounted for by activity in S1. The ability to withhold across the lockout window may be due to other wMC projections, such as to the dorsolateral striatum (Mao et al., 2011), subthalamic nucleus (Li et al., 2020), thalamus (Nakajima et al., 2019; Wimmer et al., 2015), and/or brainstem nuclei (Ioffe, 1973). Additionally, we recognize that wMC suppression does not fully equalize target-aligned and distractor-aligned stimulus selectivity (Fig. 3E,H and Fig. 4F). Thus, we suspect the involvement of additional top-down neuromodulation (Goard & Dan, 2009; Lee & Dan, 2012; Zagha & McCormick, 2014) and/or bottom-up plasticity through training (Chen et al., 2013; Chen et al., 2015). We were surprised to observe similar behavioral effects from suppressing target-aligned wMC or distractor-aligned wMC. This similarity may be due to the strong interhemispheric connections between these regions (Porter and White, 1983; Miyashita et al., 1994; Muñoz-Castañeda et al., 2021).

When are top-down cortical inputs activated to modulate sensory processing? A widely accepted framework is that stimulus-triggered bottom-up inputs are required to activate higher order regions, which in turn deliver feedback signals (Angelucci et al., 2002; Hupé et al., 1998; Manita et al., 2015; Nassi et al., 2013). In this ‘reactive’ framework, the bottom-up and top-down signals in sensory cortex occur sequentially and are temporally dissociable (Cauller, 1995; Cauller & Kulics, 1988; Kulics, 1982; Kulics et al., 1977; Manita et al., 2015). Additionally, this framework provides little motivation for studying the impacts of top-down regions on pre-stimulus activity. However, we casually demonstrated that wMC robustly modulates S1 before stimulus onset in both anesthetized and awake behaving animals (see also Jung et al., 2022). Importantly, this pre-stimulus modulation is context-dependent, with a stronger drive onto putative inhibitory neurons in distractor-aligned S1. Consistent with pre-stimulus modulation, changes in S1 sensory processing during wMC suppression are evident early, during the initial bottom-up sensory phase. These findings are more consistent with a ‘proactive’ framework, in which top-down regions sets the initial conditions for sensory processing according to goal-direction and internal state well before a stimulus arrives (Zagha, 2020).

## Methods

All experimental protocols were approved by the Institutional Animal Care and Use Committee of University of California, Riverside. Both male and female wild-type (C57BL/6) (1 male and 1 female) and transgenic (VGAT-ChR2-YFP) (7 male and 3 female) mice were purchased from Jackson Laboratories and were subsequently housed and bred in an onsite vivarium. The mice were kept in a 12/12 h light/dark cycle, and the experiments were conducted during the light phase. At the beginning of the experiments, all mice were between 1.8 and 4.2 months old (age: 85.7±8.2 days).

### Surgery

Mice were anesthetized using an induction of ketamine (100 mg/kg) and xylazine (10 mg/kg) and maintained under isoflurane (1-2% in O_2_) anesthesia. A lightweight metal headpost was attached to the exposed the skull over dorsal cortex, creating an 8 × 8 mm central window. Mice were treated with meloxicam (0.3 mg/kg) and enrofloxacin (5 mg/kg) on the day of the surgery and for two additional days after the surgery. For mice that were used for anesthetized recordings, the recordings were conducted immediately after surgery. For mice that were trained in the behavioral task, water restriction was initiated after recovery from surgery for a minimum of 3 days. For mice prepared for optogenetic experiments, the bone over the bilateral S1 and wMC was thinned under isoflurane anesthesia 2-3 days prior to the optogenetic perturbation. Between recordings, the skull was protected with Kwik-Cast (World Precision Instruments). For further surgical details, see Aruljothi et al., 2020 and Zareian et al., 2021.

### Behavior and whisker stimulation

MATLAB software and Arduino boards were used to control the behavioral task. Head-fixed mice were situated in the setup during behavioral sessions. Two piezoelectric benders with attached paddles were used: one placed within one whisker field and assigned as the target; the other placed within the opposite whisker field and assigned as distractor. Both paddles targeted the D2/E2-D3/E3 whiskers within their respective whisker fields and the assignments of target and distractor remained consistent for each mouse throughout training and recording. Each whisker stimulus was a single rapid deflection, with a triangular waveform of fixed velocity and variable duration/amplitude. In each recording session, two stimulus amplitudes were applied: the large amplitude was near the saturation of the psychometric curve and the small amplitude was 1/4^th^ of the duration/amplitude of the large amplitude stimulus, within the dynamic psychometric range. The amplitudes were customized for each mouse, in expert mice ranging from 1.4 ms to 11.2 ms duration. In all behavioral sessions, target and distractor amplitudes were equivalent. The number of target vs distractor trials and large amplitude vs small amplitude trials were approximately balanced and pseudo-randomly presented. Additionally, 30% of non-reward catch trials without whisker stimuli were distributed pseudo-randomly throughout the session.

A 200 ms lockout period (delay) was introduced after stimulus onset and before the response window. Licking during the lockout period was punished by aborting the current trial and resetting the inter-trial interval (6-10.5 s). After the lockout period, there was a 1-s response window. Licking during the response window of a target trial was considered a ‘hit’ and was rewarded with a fluid reward (approximately 5 μl of water) from the central lickport. All other licking was punished with resetting the inter-trial interval. No licking during the response window of a distractor trial or catch trial was considered a ‘correct rejection’ or ‘correct withholding’, respectively, and was rewarded with a shortened inter-trial interval (1.4-3.1 s) and subsequent target trial. Behavioral sessions typically lasted for 1 to 2 hours and generated 200-400 trials.

Mouse weights were maintained above 85% of their pre-surgery weights throughout the training and recording sessions. Mice were considered as experts once their discrimination of target and distractor stimuli (discrimination d prime = Z_hit rate_ – Z_false alarm rate_, in which Z is the inverse of the normal cumulative distribution function) reached a threshold (discrimination d prime>1) for three consecutive days. For all behavioral-physiological data presented in this study, mice were performing at expert level during data collection. For more training details and learning trajectories, see Aruljothi et al., 2020; for more training setup information, see Zareian et al., 2021.

### Electrophysiological recording and whisker receptive field mapping

All electrophysiological recordings were obtained from 9 VGAT mice and 2 wild type mice. On average, 7 sessions were recorded from each mouse (range 1-18). For electrode implantation, a small craniotomy with durotomy (approximately 0.5 mm in diameter) above the S1 barrel field (from bregma: 2.3-4 mm lateral, 1-2.2 mm posterior) was made under isoflurane anesthesia the same day of recording (used for up to three days of acute recordings). After approximately 30 minutes post-surgery, mice were tested in the behavioral task without electrode implantation to ensure recovery to normal behavior. Upon evidence of normal expert behavior, a silicon probe (Neuronexus A1x16-Poly2-5 mm-50s-177) was advanced into the brain using a Narishige micro-manipulator under stereoscope guidance. This silicon probe design has 16 electrode sites, arranged in two columns spanning 375 μm, allowing for tetrode clustering of single units. We positioned the recording sites to target layer 5, from 500 to 1000 um below the pial surface (mid-point of the silicon probe recording sites: 626.0 ± 10.5 μm). Whisker alignment for S1 recordings was verified by two methods (as also described in Zareian et al., 2021). First, during electrode implantation we verified correct alignment by hand mapping of several individual whiskers and observing real-time local field potential (LFP) display. Second, post-hoc we plotted the large amplitude stimulus evoked multi-unit spiking activity and only included sessions with robust and fast-rising peaks in the post-stimulus time window (peak response reaching at least 1.4×baseline activity within a 50 ms post-stimulus window).

### Optogenetic suppression and opto-tagging

To transiently suppress specific cortical regions, we used optogenetic stimulation of GABAergic interneurons in VGAT-ChR2-YFP mice (Guo et al., 2014; Zhao et al., 2011). Control studies with identical optical illumination were performed in wild type mice without ChR2 expression. For optogenetic stimulation, LED fiber optic terminals were placed above wMC (centered on [from bregma]: 1 mm lateral, 1 mm posterior) and/or S1 (same coordinates as above). Optogenetic GABAergic stimulation was used for both suppression studies as well as for opto-tagging of recorded GABAergic interneurons. For both applications, 470 nm blue light was delivered through a 400 µm core diameter optical fiber (Thorlabs). Optogenetic stimulation was delivered using a triangular function, beginning 200-500 ms before whisker stimulus onset, peaking 100-200 ms after stimulus, and decaying for 400-600 ms. This protocol was used to 1) prevent transient onset/offset effects of square pulse illumination and 2) ensure a relatively strong and stable suppression for 100 ms pre-stimulus through 100 ms post-stimulus. Maximum intensity at peak stimulation was 18 mW/mm^2^ (2.26 mW) at the fiber tip. For the behavioral sessions without electrophysiological recordings, optical suppression was applied on one third of all trial types, randomly distributed. For the sessions (awake or under anesthesia) with electrophysiological recordings, simultaneous optical suppression and opto-tagging was conducted, in which optical suppression was randomly applied on one third of all trial types and opto-tagging was randomly applied on 7-10% of all trial types.

Barrier and masking methods were used to prevent visual detection of the optical stimulus. Barrier methods consisted of wrapping the headpost and optical fiber in opaque material. Masking consisted of applying a 10 Hz blue light LED stimulation (duty ratio=50%) directly above the subject’s eyes on all trials, beginning 180 ms before optical stimulation onset and ending 170 ms after optical stimulation offset. Masking light was given on all trials (irrespective of suppression or stimulus type).

### Electrophysiological recording data pre-processing

Neuralynx software was used for data acquisition and spike sorting. Putative spikes were identified as threshold crossings over 18-40 μV. Spike sorting and clustering was conducted offline by SpikeSort3D software, first through the KlustaKwik function followed by manual inspection of waveform and inter-spike interval distributions. On average, 23 units were identified in each recording session (22.8±5.4). Further data analyses were conducted using MATLAB software (MathWorks). Spike times were binned within 5 ms non-overlapping bins. For more details about spike sorting, see Zareian et al., 2021.

### Widefield Ca^2+^-sensor imaging

Neuronal activity of dorsal cortex was imaged in GCaMP6s expressing mice during expert performance in the selective detection task. The imaging datasets analyzed here were previously published (Aruljothi et al., 2020; Marrero et al., 2022). The dataset consists of n=40 behavioral-imaging sessions from n=5 mice. Analyses included all target trials and all distractor trials, regardless of behavioral outcome. Imaging data were processed for pixel-by-pixel stimulus encoding (neurometric d prime), comparing pre-stimulus (stimulus absent) and post-stimulus (stimulus present) time points (Aruljothi et al., 2020). The analyses presented in this study are from the 100-200 ms post-stimulus imaging frame, during the lockout period and before the response window. Regions of interest (ROI) were selected to include dorsal primary somatosensory cortex, including the dorsal portion of the whisker field. Analyses focused on stimulus encoding in the unpreferred hemisphere (ipsilateral to the whisker stimulus). D prime values were averaged within each ROI and compared for target and distractor stimulus trials.

### Data analysis

All data were analyzed using SPSS or custom-written MATLAB scripts. ANOVA was conducted in SPSS by univariate general linear model. Data are reported as mean +/- the standard error of the mean (SEM), unless otherwise stated.

Behavioral detection d prime: d’= Z_Hit Rate_ – Z_Spontaneous Licking Rate_. Z is the inverse of the normal cumulative distribution function. Response rates of 0 and 1 were adjusted as 0.01 and 0.99 respectively to avoid yielding infinite values. Neuronal encoding d prime across a 100 ms post stimulus window: d’=√2×Z_AU-ROC._ AU-ROC is the area under the receiver operating curve (ROC), in which the curve is composed of the spike counts within 100 ms pre-stimulus windows and 100 ms post-stimulus windows. Each window was composed to two 50 ms epochs. Neuronal encoding d prime across 20 ms post stimulus sliding windows: d’=√2×Z_AU-ROC._ This calculation is similar to that described above, except the post stimulus window was a single 20 ms window and was compared to 5 consecutive 20 ms pre-stimulus windows. The post-stimulus 20 ms sliding window was sampled every 5 ms, thereby consisting of 75% overlap, up to 100 ms post-stimulus. To account for the 5x sampling of the pre-stimulus window, we multiplied the post-stimulus histogram by 5 before plotting the ROC. AU-ROC values of 0 and 1 were adjusted as 0.003 and 0.997 to avoid yielding infinite values.

For the linear fitting of the single unit population data and to determine the slopes with 95% confidence intervals, we used MATLAB functions: the polyfit function was used to calculate the slope for the middle linear fitted line, and the polyparci function was used to calculate the 95% confidence intervals. These data were plotted by the fitlm function.

Classification analyses: We trained a linear discriminant model (MATLAB function: fitcdiscr, with diagonal covariance matrix) to classify the spike rates of target-aligned S1 neurons into target or distractor trial types. Out of all the putative excitatory units recorded from target-aligned S1 (n=197), the half of units with largest target stimulus encoding d prime (n=98) were pooled together as pseudo-ensembles. Spike rates on each trial were determined as the increase in spiking from pre-stimulus baseline, using only the first 50 ms post-stimulus in order to better isolate the initial sensory response. To equalize the number of trials across sessions, we reconstructed the data: the sessions with less trials had their trials duplicated and appended to the original trials to match the trial number of the session with the most trials; for the sessions with the trial numbers not a common divisor of the trial number in the longest session, the trials were randomly sampled (without replacement) from these sessions accordingly and added to that session to fill in. After data reconstructions, each unit contained 70 target or distractor light-off trials and 30 target or distractor light-on trials. For each simulation, the trial order for all the units were randomly shuffled to remove the temporal structure and correlation between units. 40 target light-off trials and 40 distractor light-off trials were used to train the linear discriminant model. We then tested the model on the 30 light-off target and distractor hold-out trials and the 30 light-on target and distractor wMC suppression trials. This process was iterated 1000 times. Data (Supplementary Figure 7) are presented as mean ± standard deviation.

### Data inclusion criteria and quality control

‘Task engaged’ was defined as a continuous period of at least 10 min with no greater than a 60 s gap of non-licking. For any session with more than one engaged period, only the longest continuous segment was used for further analyses. Sessions without continuous engagement for at least 10 min were excluded from further analysis. ‘Task disengaged’ epochs always followed periods of task engagement. Disengaged epochs were recorded for approximately 30 min and contained at least 300 trials. We suspect that task disengagement was due to lack of motivation from satiety after receiving sufficient rewards.

For analyses of behavioral outcomes, at least 5 trials of a given type were required to be included in the session analysis. For calculations of spontaneous licking rate, we used the 1s pre-stimulus window of all trial types. These values were similar to the spontaneous licking rates based on catch trials yet included substantially more trials in their calculation.

For recording data indicated as behaving/target/distractor sessions, we analyzed only the ‘task engaged’ epochs as defined above, so that the behavioral and neuronal data were based on the same trials. In Fig. 5 B, ‘VGAT’ is based on all sessions recorded from VGAT-ChR2 mice, regardless of the engagement status or wakefulness. Similarly, Supplementary Fig. 5 is based on all recording sessions.

#### Principal whisker alignment inclusion criteria

Out of 78 recording sessions, 14 were unaligned, 64 were aligned, based on criteria described above. Only the aligned sessions are used for neuronal data analyses. wMC suppression sessions with unaligned S1 recordings were included for behavior only analyses, along with wMC suppression sessions without extracellular recordings.

#### Visual mask-induced noise filtering

For optogenetic sessions, we applied a 10 Hz mask light above the eyes of mice, which sometimes caused a 10 Hz (onset triggered) or 20 Hz (onset and offset triggered) signal in the neuronal recording data (likely due to visual transduction). To minimize the influence of this contamination, we applied a band-pass filters (‘designfilt’ function of MATLAB, designed based on Butterworth notch filter) to the spike count time series for single unit or multi-unit data before performing subsequent analyses.

#### Putative excitatory and putative inhibitory neurons inclusion criteria

To identify putative excitatory or inhibitory units, we analyzed the time window of the optogenetic light duration (excluding the first and last 100 ms time bins at the start and the end of the triangle waveform) to calculate the averaged trial-by-trial light-on spike rate for each unit (using an equivalent window for light-off control trails). We calculated the difference of averaged rates between light-on trials and light-off trials, and determined significance from two-sample t tests with a threshold of p<0.1. Out of these significantly modulated units, if the average spike rate of light-on trials was larger than that of light-off trials (enhanced), the units were considered as putative inhibitory neurons; if the average spike rate of light-on trials was smaller than that of light-off trials (suppressed), the units were considered as putative excitatory neurons. Any unit with p>=0.1 was not categorized and not included in further cell-type analyses.

## Acknowledgments

We thank the members of the Zagha lab for discussions regarding experimental design and data analyses. We especially thank Krithiga Aruljothi and Krista Marrero for providing the widefield imaging dataset. We also thank Dr. Hongdian Yang and Dr. Chunyu Ann Duan for their valuable comments on a previous version of this manuscript. This research project was supported by NIH/NNDS R01 Grant NS107599 and Whitehall Foundation Grant 2017-05-71.

## Author Contributions

Z.Z. and E.Z. initiated the project and designed the experiments. Z.Z. performed the experiments and acquired the data. Z.Z and E.Z. analyzed the data and prepared the manuscript.

## Supplementary material

**Supplementary Fig. 1:**
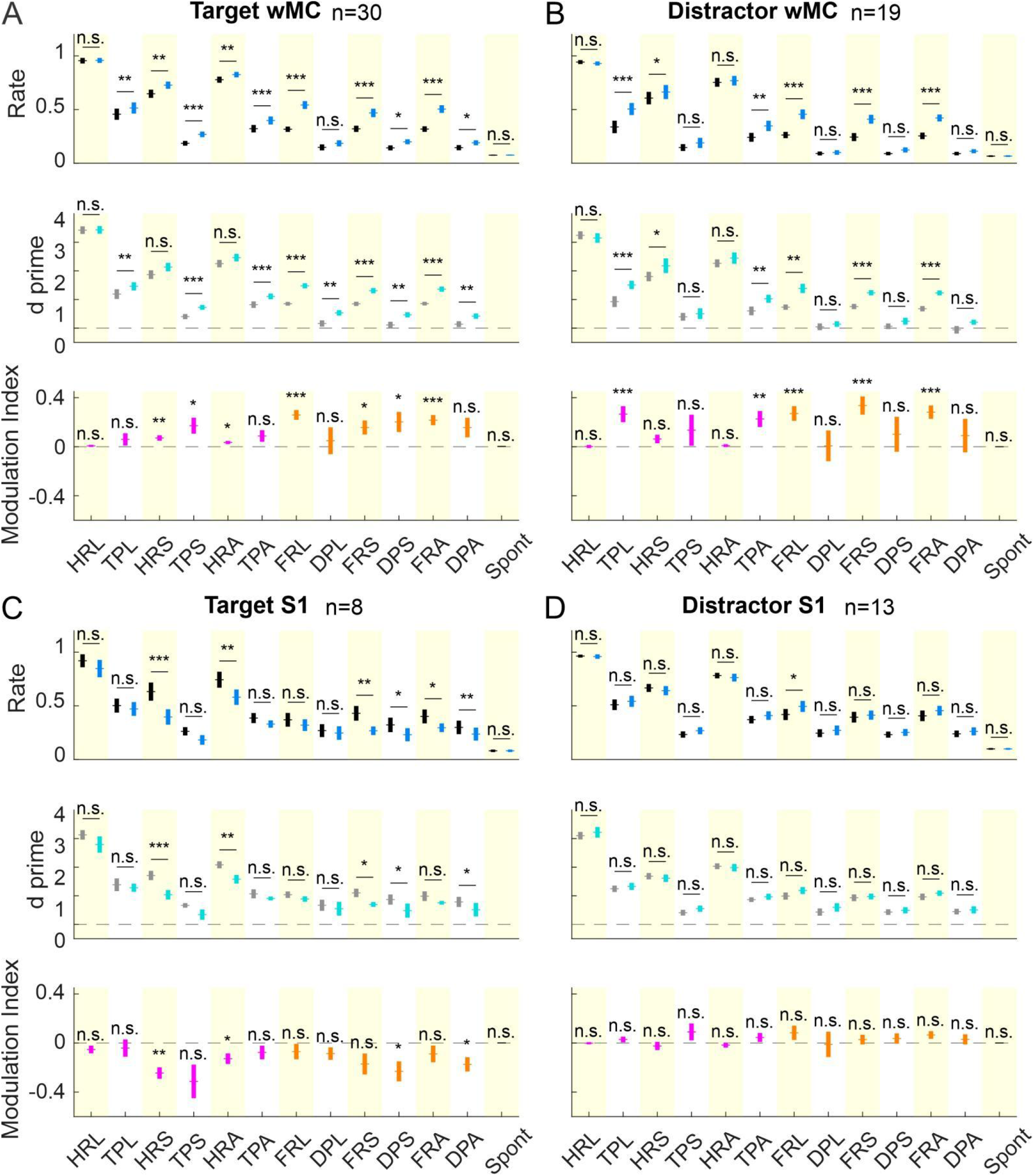
Behavioral effects of optogenetically suppressing target-aligned wMC (A), distractor-aligned wMC (B), target-aligned S1 (C), and distractor-aligned S1 (D). HRL: hit rate for large amplitude stimuli; TPL: target premature licking rate for large amplitude stimuli; HRS: hit rate for small amplitude stimuli; TPS: target premature licking rate for small amplitude stimuli; HRA: hit rate for all stimulus amplitudes (large and small); TPA: target premature licking rate for all stimulus amplitudes; FRL: false alarm rate for large amplitude stimuli; DPL: distractor premature licking rate for large amplitude stimuli; FRS: false alarm rate for small amplitude stimuli; DPS: distractor premature licking rate for small amplitude stimuli; FRA: false alarm rate for all stimulus amplitudes; DPA: distractor premature licking rate for all stimulus amplitudes; Spont: spontaneous licking rate. Control (light-off) data are in black; wMC optogenetic suppression (light-on) data are in blue. Yellow shaded areas separate non-premature rates (shaded) from premature rates (not shaded). See Methods for calculations of response rates (top row), d prime (stimulus response rates normalized by spontaneous response rates, middle row), and modulation index (comparison between control and wMC optogenetic suppression trials, bottom row). As in the rest of the study, p values are indicated as: n.s. (p>=0.05), *(0.01<=p<0.05), **(0.001<=p<0.01), or ***(p<0.001).

**Supplementary Fig. 2:**
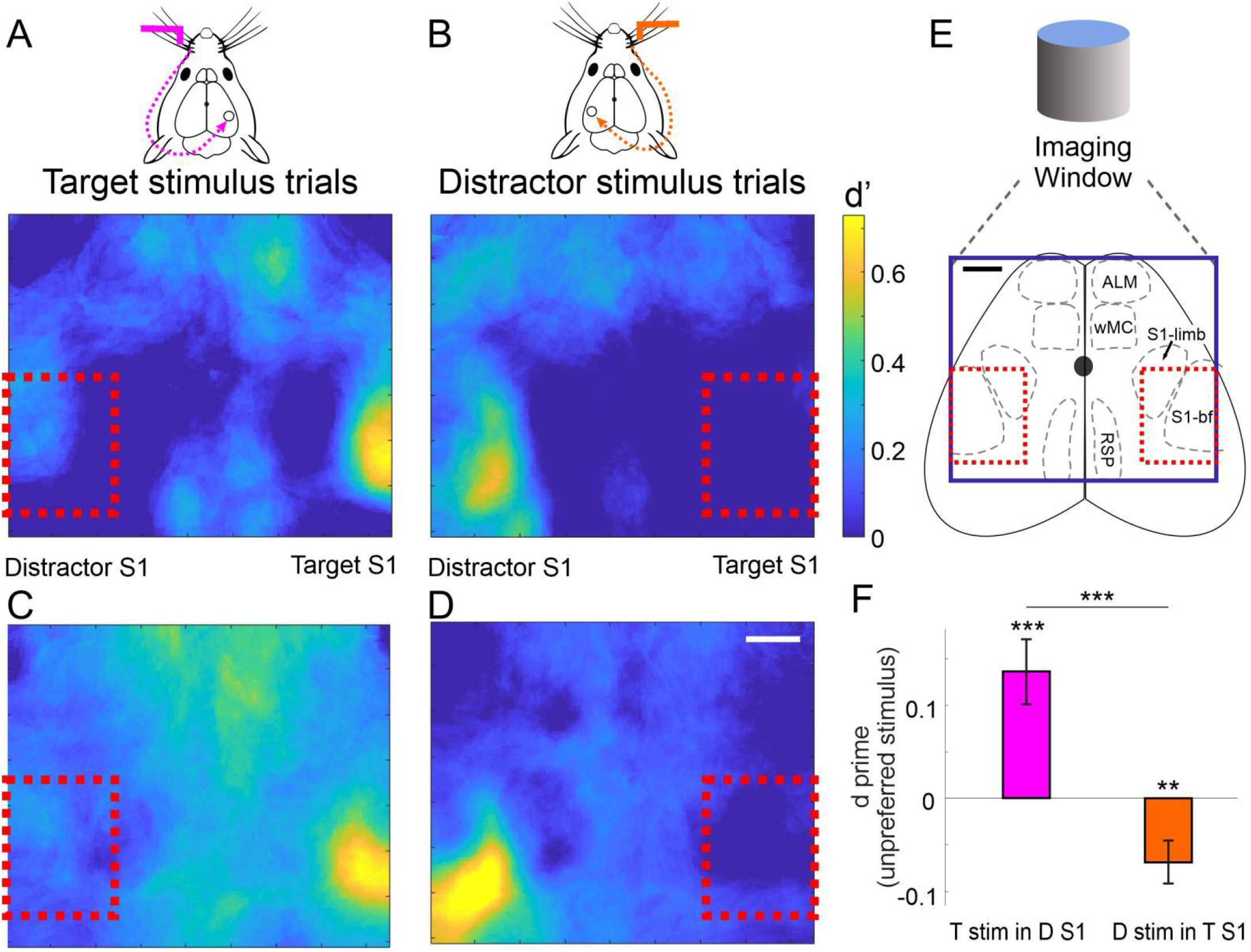
Widefield GCaMP6 imaging evidence for asymmetric activation of target-aligned and distractor-aligned S1 in response to their unpreferred stimuli. (E) A reference brain map, showing the imaging window presented in [A-D], with a few task-relevant regions indicated. ALM: anterior lateral motor cortex; S1-limb: primary somatosensory cortex, limb regions; S1-bf: primary somatosensory cortex, barrel field; RSP: retrosplenial cortex. The analysis regions of interest (ROIs) are outlined in red. (A-D) Data from two example imaging sessions. Stimulus encoding (d’) was computed for each pixel as the separation between stimulus absent and stimulus present, for target stimulus trials [A,C] and distractor stimulus trials [B,D]. Activations evoked by unpreferred stimuli were quantified (target stimuli in distractor-aligned S1 and distractor stimuli in target-aligned S1). (F) Quantification of averaged stimulus encoding (d prime) in the selected ROIs calculated across n=40 sessions. Above each bar are results from one-sample t tests to determine differences from zero within each ROI, and the line between bars reflects results from a two-sample t tests to determine differences between ROIs. T stim in D S1, target stimulus encoding in distractor-aligned S1; D stim in T S1, distractor stimulus encoding in target-aligned S1. Scale bars in D, E, 1 mm.

**Supplementary Fig. 3:**
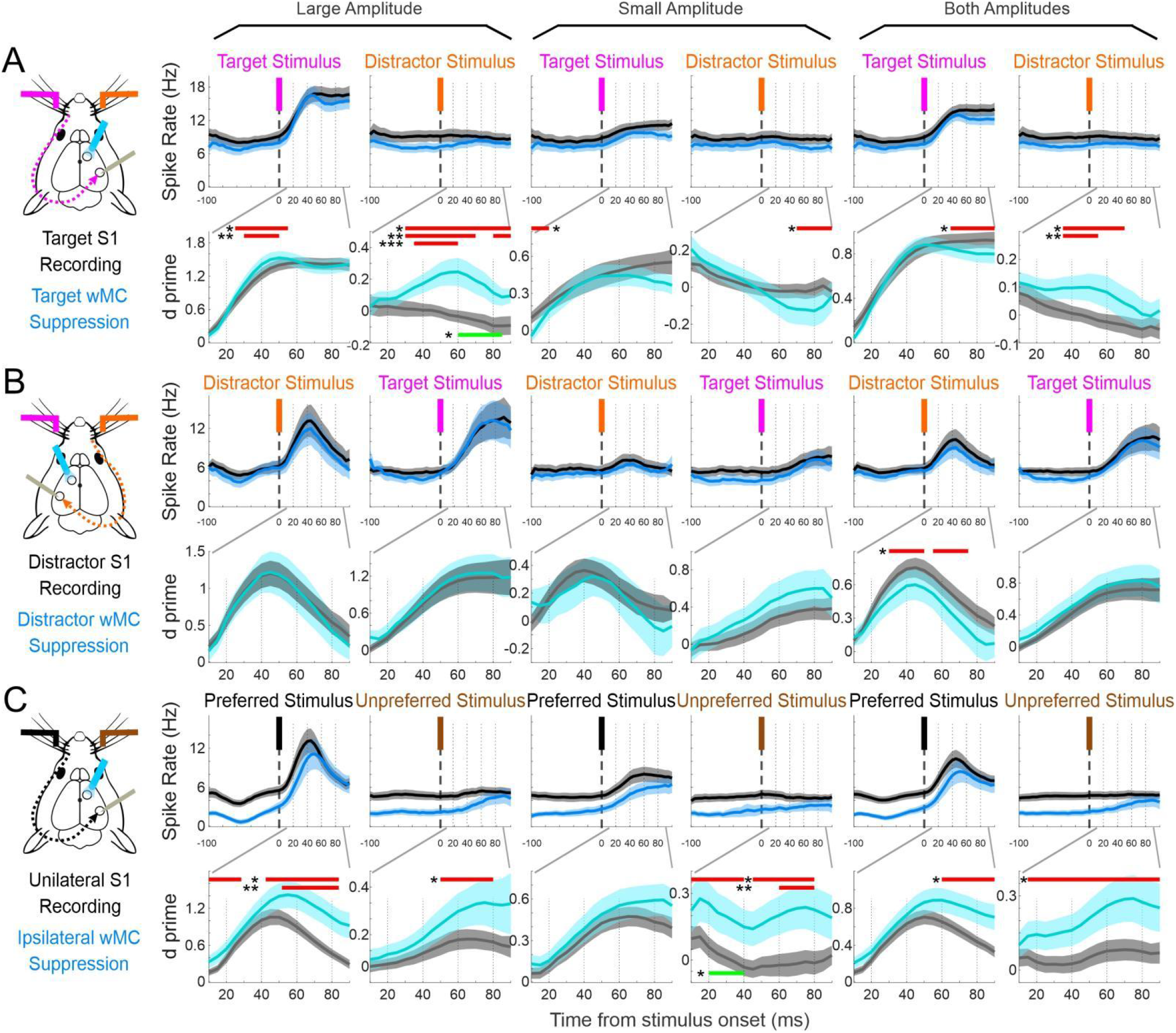
Neuronal spike rates and d prime of S1 neurons during wMC suppression in response to different stimulus amplitudes (following the same plotting as Fig. 3C-H). (A) Target-aligned S1 recordings and target-aligned wMC suppression. (Top) Black traces indicate averaged spike rates across all sessions during light-off control trials while dark blue traces indicate light-on wMC suppression trials. (Bottom) Gray traces indicate neuronal d prime across all session from light-off control trials while light blue traces indicate light-on wMC suppression trials. (B) Same as [A] but for distractor-aligned S1 recordings and distractor-aligned wMC suppression. (C) Same as [A] but for naïve mice recorded under anesthesia. In all plots, curves indicate average values ± standard error (shaded). The red bars indicate d prime value pairs that are significantly different (paired t test); green bars indicate d prime values from light-off control conditions are significantly lower than zero (one sample t test).

**Supplementary Fig. 4:**
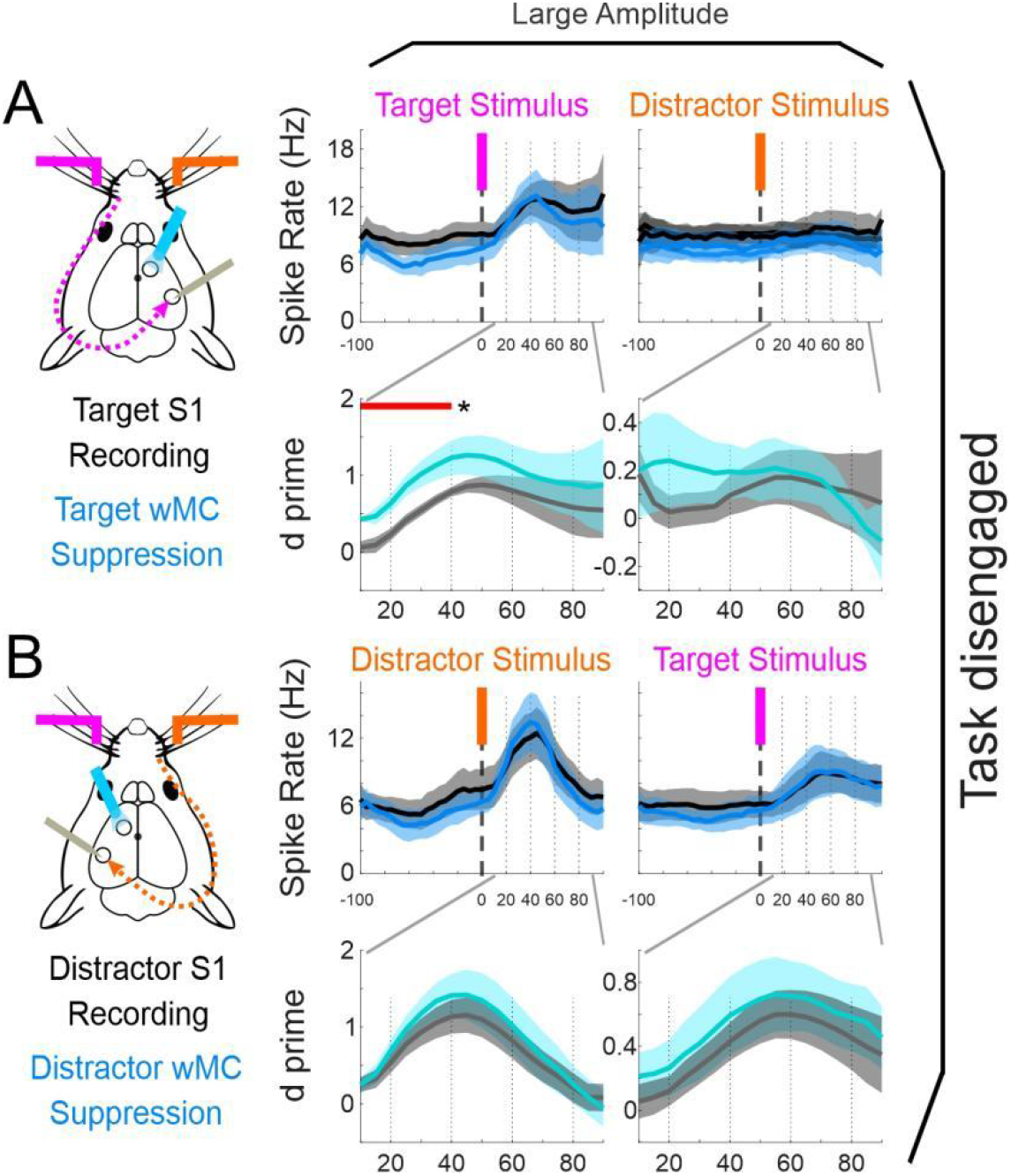
Effects of wMC suppression in task disengaged mice. Data are presented as in Fig. 3C-H. (A) Target-aligned S1 recordings and target-aligned wMC suppression during disengaged recording sessions. (B) Distractor-aligned S1 recordings and distractor-aligned wMC suppression during disengaged recording sessions. wMC suppression caused minimal effects on stimulus encoding during task disengagement.

**Supplementary Fig. 5:**
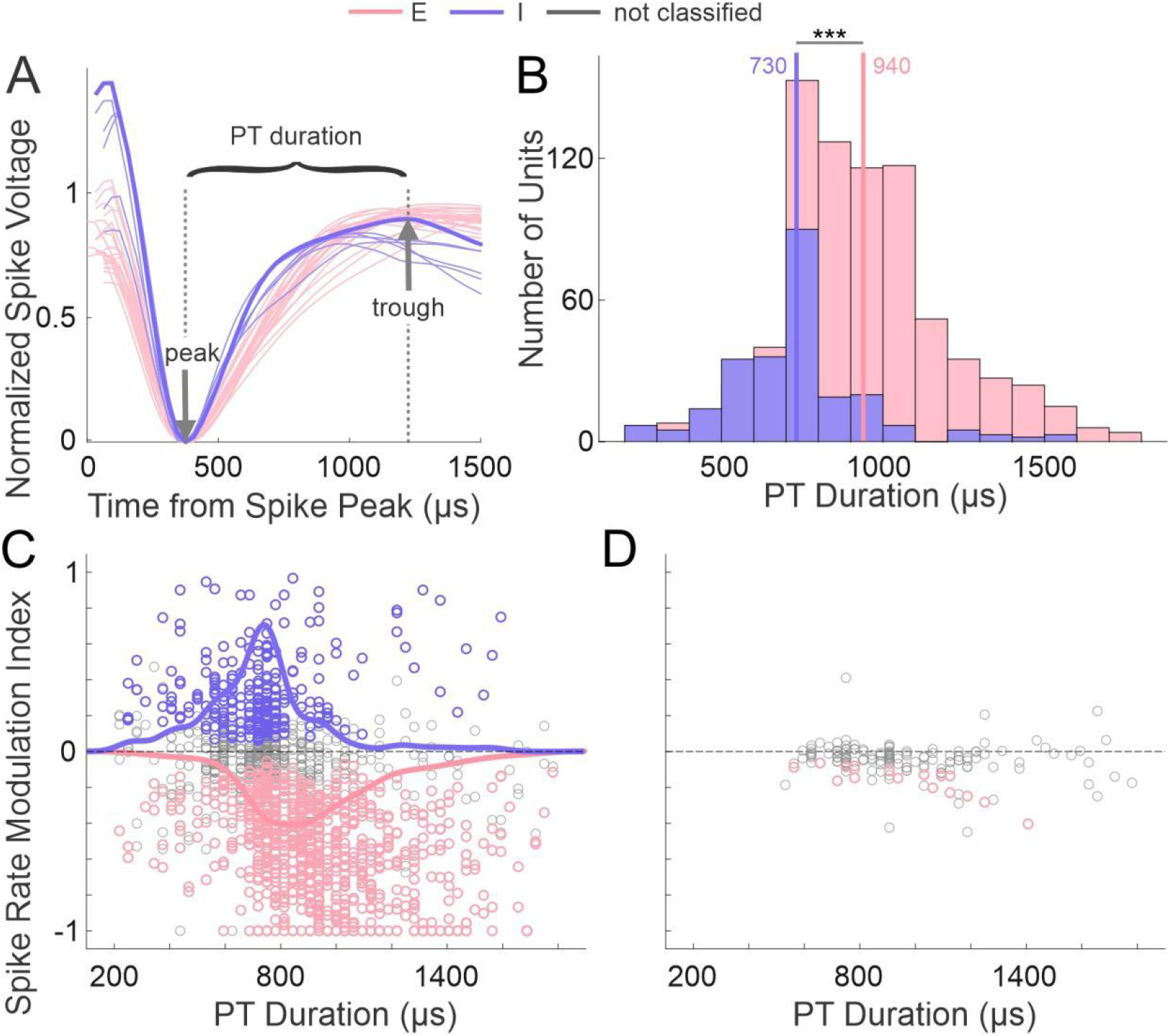
Opto-tagging method and verification. Putative excitatory or putative inhibitory neurons are assigned based on significant decreases or increases, respectively, in spike rate during opto-stimulation within the S1 recording site, using p<=0.1 as the significance threshold. (A) Spike widths of putative inhibitory (purple) and excitatory (pink) neurons from one example session. Spike width is calculated as peak-trough (PT) duration. (B) Histogram of putative excitatory and inhibitory neuron counts of different spike widths. Two vertical lines indicate the average spike widths of each population, which are significantly different from each other (two sample t test, p=1.73×10^-28^). Putative inhibitory neurons have shorter spike widths, as expected, yet there is substantial overlap between these populations. (C) Distribution of spike width versus opto-tagging modulation index from VGAT-ChR2 mice (n=72 recording sessions). Each data point is one unit, and the two curves show the density distributions. Modulation index (MI) is calculated as the change in spike rate during opto-tagging. MI>0 indicates units enhanced by direct light stimulation; MI<0 indicates units suppressed by direct light stimulation. 1613 neurons are recorded, in which 762 neurons (47.24%) are characterized as putative excitatory neurons, 246 neurons (15.25%) are characterized as putative inhibitory neurons, 605 neurons (37.51%) are not significantly modulated by opto-tagging and therefore unassigned. (D) As same as [C] but the units were recorded from control WT mice (n=6 recording sessions). In WT mice, 155 neurons were recorded, in which 16 neurons (10.03%) are significantly modulated by opto-tagging, consistent with an expected false positive rate of 10%.

**Supplementary Fig. 6:**
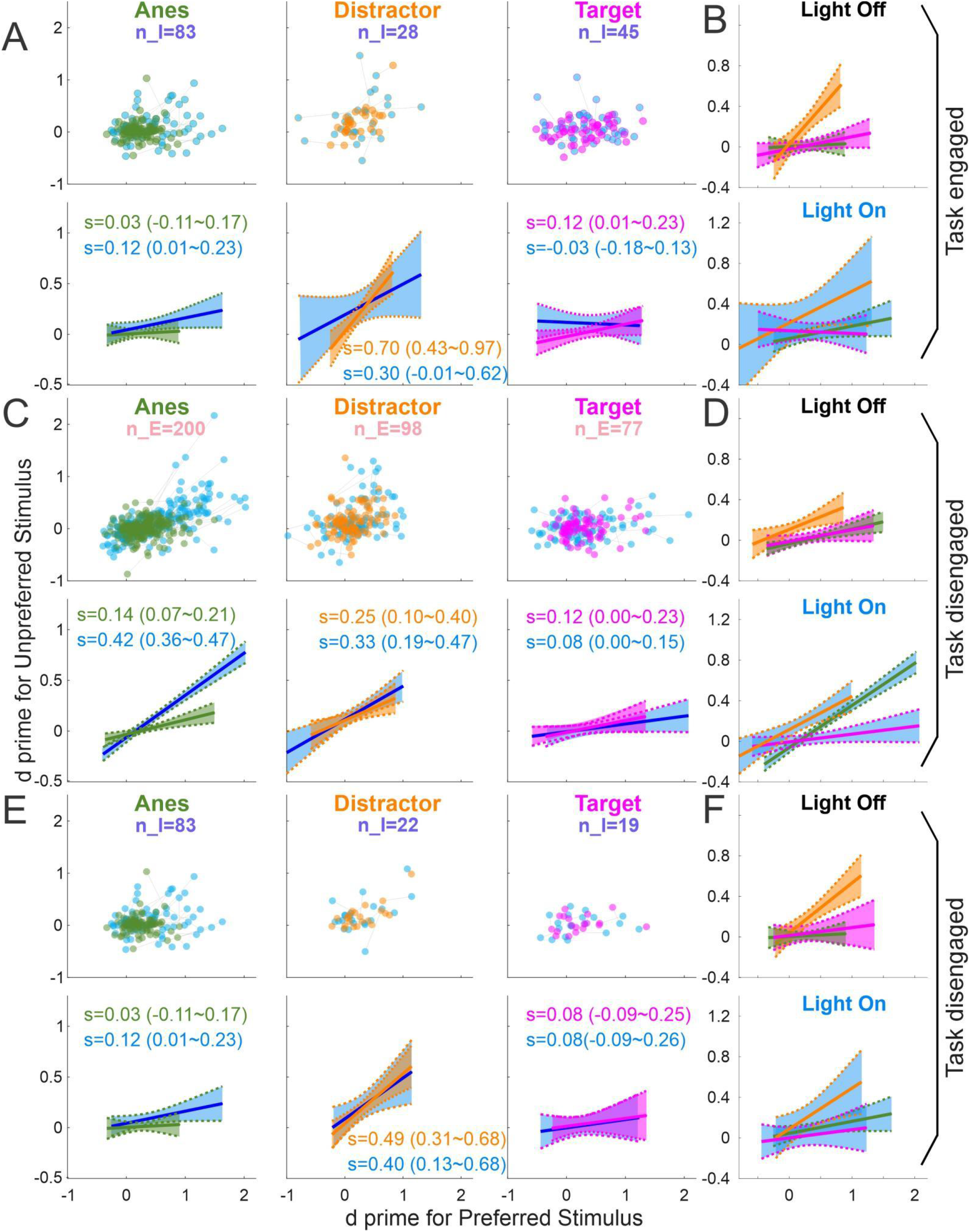
S1 neuronal encoding of preferred and unpreferred stimuli, according to cell type and task engagement. (A) As same as Fig. 4 D-F, but for putative inhibitory neurons recorded: under anesthesia (1^st^ column), in distractor-aligned S1 in awake behaving mice (2^nd^ column), and in target-aligned S1 in awake behaving mice (3^rd^ column). (B) The overlay of the control (top) and wMC suppression (bottom) linear fits from the second row of [A]. (C, D) As same as Fig. 4 [D-G] (putative excitatory neurons), but for disengaged awake recording sessions. (E, F) As same as Fig. 4 [D-G], but for inhibitory neurons recorded from disengaged awake recording sessions. Anesthetized data is duplicated for ‘engaged’ and ‘disengaged’ categories and included for reference. Note, for putative excitatory neurons when task disengaged [C], the stimulus selectivity profiles for distractor-aligned (orange) and target-aligned (magenta) S1 neurons are not significantly different. Furthermore, for these conditions, wMC suppression (blue) does not change stimulus selectivity. These data are in marked contrast to S1 excitatory neurons during task engagement (Fig. 4 E,F).

**Supplementary Fig. 7:**
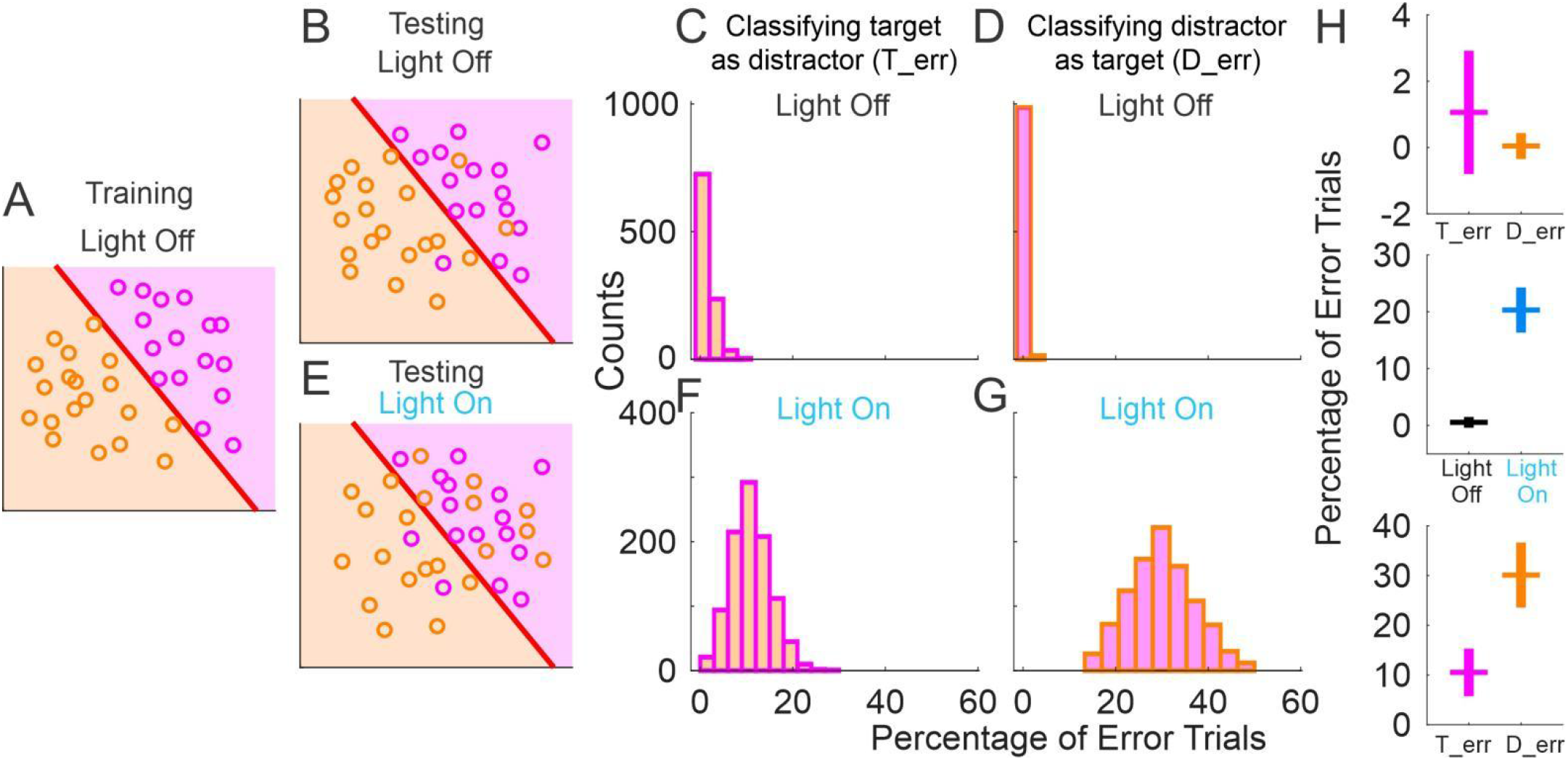
Classification outcomes from linear discriminant analyses of trial types. (A) Illustration of a hypothetical linear model separating trial types based on training data. Each circle indicates one trial. Magenta circles indicate true target trials; orange circles indicate true distractor trials. The line indicates the classifier. In this example, the model correctly classifies all target and distractor trials. (B) Illustration of classification of hold-out control trials. (C,D) Classification data from hold-out control trials. Histograms of percent error bins for miss (C, classifying target as distractor) and false alarm (D, classifying distractor as target) errors, from 1000 iterations. (E) Illustration of classification of wMC suppression trials. (F,G) Classification data from light-on wMC suppression trials, plotted as in [C, D]. (H) Mean ± standard deviation comparisons for [C] vs [D] (top,T_err %: 1.06±1.86; D_err %: 0.05±0.39), [F] vs [G] (bottom, T_err %: 10.52±4.79; D_err %: 30.09±6.56), and the average of [C] and [D] vs the average of [F] and [G] (middle, Light-off %: 0.55±0.95, Light-on %: 20.31±3.98).

## Notes

### Competing Interest Statement

The authors have declared no competing interest.

### Summary of Updates

A new figure is added between original Fig1 and Fig2; A new supplementary is added between original supplementary Fig1 and supplementary Fig2.

